# OPA1 controls mitochondrial dysfunction-driven liver fibrosis in MASLD

**DOI:** 10.64898/2026.09.24.753666

**Authors:** Xinting Yu, Pinzhu Huang, Arvo Justice, Aaron Hakim, Yoshiko Nanki, Zaur Abilov, Edward B. O’Neill, Jerry Cruz Rodriguez, Enzo Cabrera, Marla Sagatelian, Chandra S. Shankar, Brenda R. Flam, Kawtar Al Khalloufi, Hiromi Sesaki, Shisong Rong, Yury V. Popov

**Affiliations:** Department of Internal Medicine, University of South Florida, Tampa, FL; Department of Medicine, Division of Gastroenterology, Hepatology & Nutrition, Beth Israel Deaconess Medical Center, Harvard Medical School, Boston, MA; Broad Institute of MIT and Harvard, Cambridge, MA; Department of Pathology and Cell Biology, University of South Florida, Tampa, FL; Department of Gastroenterology and Hepatology, Liver Transplant, Tampa General Hospital, University of South Florida, Tampa, FL; Department of Cell Biology, Johns Hopkins University School of Medicine, Baltimore, MD, USA; Ocular Genomics Institute, Mass Eye and Ear, Harvard Medical School, Mass General Brigham, Boston, MA

**Author notes:** **Correspondence to:** Yury V. Popov MD, PhD, Division of Gastroenterology, Hepatology and Nutrition, Beth Israel Deaconess Medical Center, Harvard Medical School, 99 Brookline Ave., RN-234, or: Shisong Rong MD, PhD, Department of Ophthalmology, Ocular Genomics Institute, Mass Eye and Ear, Mass General, Brigham, Harvard Medical School, Boston, MA, United States, 243 Charles St., Boston, 02114 MA, USA.

## Abstract

Progressive hepatic fibrosis is the principal determinant of morbidity and mortality in metabolic dysfunction-associated steatotic liver disease and steatohepatitis (MASLD/MASH). Mitochondrial dysfunction is a hallmark of MASH, and the release of mitochondrial damage-associated molecular patterns (mito-DAMPs) from injured hepatocytes can promote fibrosis. However, how mitochondrial dynamics and quality control shape the fibrotic response in MASLD/MASH remains unclear. Here, through large-scale genomic analyses of mitochondrial genes governing mitophagy, fusion and fission in human MASLD, with a power-equivalent sample size of approximately 700,000 individuals, we identify a strong association between hepatic fibrosis and the mitochondrial fusion factor dynamin-like GTPase optic atrophy 1 (*OPA1*). *OPA1* transcripts and protein abundance in the liver epithelium were progressively dysregulated with advancing fibrosis. In mice, hepatocyte-specific *OPA1* loss alone was sufficient to induce hepatic stellate cell activation and fibrosis in zone 3, promoted the release of mito-DAMPs into the circulation and exacerbated fibrosis in experimental MASH. These findings identify *OPA1* as a central regulator of the hepatic fibrotic response and connect defective mitochondrial homeostasis to mito-DAMP release, hepatic stellate cell activation and fibrosis in MASLD.

## Introduction

Metabolic dysfunction–associated steatotic liver disease (MASLD) has emerged as the most prevalent chronic liver disease worldwide, tracking closely with the global epidemics of obesity and type 2 diabetes.(1) Current estimates place adult MASLD prevalence at approximately 30%-40% globally, 60%-70% in type 2 diabetes, and 70%-80% in obesity, making it a leading driver of liver-related morbidity and a major contributor to global health-care utilization.(1, 2) The recently adopted steatotic liver disease (SLD) nomenclature, including MASLD and metabolic dysfunction–associated steatohepatitis (MASH), reflects the central role of cardiometabolic risk factors in disease pathogenesis and provides a more inclusive, pathophysiology-aligned framework for research and clinical care.(3, 4) Epidemiologic syntheses and updated disease primers underscore that MASLD prevalence continues to rise across regions, with substantial heterogeneity driven by demographic structure, metabolic comorbidity burden, and differences in health systems.(5, 6)

Although most individuals with MASLD have benign non-progressive steatosis, about a quarter of patients develop steatohepatitis (MASH), which is characterized by low-grade liver injury, inflammation, and fibrosis.(1, 2) Fibrosis, if progresses, eventually leads to end-stage cirrhosis and life-threatening sequala e. Multiple retrospective analyses of biopsy-confirmed MASLD cohorts linked histologic fibrosis stage—rather than necroinflammatory activity—to liver-related morbidity and mortality.(7–9) Most recently, a large prospective multicenter study of 1773 patients confirmed that of all histological disease characteristics, only advanced fibrosis (CRN F3-F4) predicts an increased risk of liver-related complications and death in MASLD.(10) These studies have unequivocally established progressive fibrosis as a major driver of clinically significant MASLD, ighlighting the urgent need to define mechanistic pathways that determine why some patients transition from metabolic liver injury to progressive fibrogenesis and eventually develop end-stage liver disease. Precise understanding of underlying genetics and molecular networks is needed to develop novel, effective antifibrotic therapies for MASH.

Hepatocyte injury and innate immune activation are key prerequisites of fibrogenic responses, but the mechanisms underlying why only a subset of patients with steatohepatitis develop exaggerated, progressive fibrosis remain incompletely understood. Sterile injury triggers dying cells to release of damage-associated molecular patterns (DAMPs) into extracellular space, which alarm the immune system to mount inflammatory response via activation of pattern-recognition receptors(11). Specifically, mitochondria-derived danger signals (mito-DAMPs) released from damaged cells have recently emerged as a major source of fibrogenic DAMPs in human MASH.(12) The liver is an organ that is particularly rich in mitochondria due to its critical metabolic function in the body, with each hepatocyte containing 1000-2000 mitochondria making up to 25% of the cell volume, and each mitochondrion containing 2-10 copies of the mitochondrial genome.(13) Mito-DAMPs are considered uniquely biologically potent due to their bacterial ancestry and structural similarity. (14) Human mitochondrial DNA (mtDNA) has a circular covalently-closed structure (15) with non-methylated CpG repeats,(16) thus bearing structural resemblance to highly immunogenic bacterial DNA.(17) Other mito-DAMPs, including N-formyl peptides, cardiolipin, ATP, and cytochrome C, can also induce inflammatory responses when released into the extracellular space. (18)

Recently, we have identified mito-DAMPs released from dying hepatocytes as major direct driver of fibrotic response.(12) Exposure to mito-DAMPs in vivo or in vitro leads to rapid activation of hepatic stellate cells (HSCs), the principal fibrogenic effector cells of the liver, with mtDNA identified as major active component. Circulating mito-DAMPs are elevated in human MASH, and further increased in subset of patients with advanced fibrosis.(12) Moreover, mtDNA sensing pathways in liver macrophages have been implicated in the progression of steatohepatitis and fibrosis; for example, Stimulator of Interferon Genes (STING) signaling in Kupffer cells can act as an mtDNA-responsive axis promoting inflammatory outputs and disease severity in murine MASH models.(19) Collectively, these data support a model in which mitochondrial injury is not merely a byproduct of metabolic stress, but an upstream instigator of immune–stromal crosstalk that drives fibrogenesis.

A critical determinant of whether mitochondrial injury culminates in intra- and extra-cellular mito-DAMP escape is mitochondrial quality control (MQC).(20) MQC comprises coordinated mitochondrial fusion and fission, biogenesis, and selective autophagic removal of damaged organelles (mitophagy)—processes that preserve bioenergetic capacity while limiting oxidative stress and inflammatory leakage. Reviews of mitochondrial homeostasis in steatotic liver disease emphasize that perturbations in mitochondrial dynamics and mitophagy accompany MASLD progression and may influence both hepatocyte survival and inflammatory tone.(21, 22) Conceptually, defective MQC could create a feed-forward loop: lipid overload and oxidative injury impair mitochondrial turnover, damaged mitochondria accumulate and release mito-DAMPs, innate immune pathways intensify inflammation, and inflammatory and profibrotic programs further destabilize cellular metabolism.

We hypothesized that unknown genetically determined MQC defects in subset of patients with MASLD may underlie the escape of mito-DAMPs to trigger progressive fibrosis. Deciphering molecular machinery linking mitochondrial disfunction in MASH to pro-fibrogenic mito-DAMPs release is clinically relevant – to help identify those patients who are at risk of clinical progression to cirrhosis and to elucidate novel therapeutic targets to slow down, halt or reverse fibrotic MASH. In this study, we performed a mitochondria-focused large scale human genetics study and functional validation experiments to elucidate the mechanistic links between mitochondrial biology, mito-DAMP release and fibrotic responses, to identify molecular underpinning upstream of mito-DAMPs-driven fibrogenesis in MASH.

## Methods

### 1.1 Study workflow and design of the analytical pipeline

Figure 1 summarized the study workflow. Briefly, first we performed an integrative genetic analysis of common and rare variants to evaluate whether genes involved in mitochondrial dynamics and MQC are associated with MASLD and liver fibrosis-related traits. This followed by functional annotation using expression quantitative trait locus data(23) and publicly available tissue gene expression databases. (24–26) Identified candidate gene product expression was studied in patient-derived MASH liver tissue at different fibrosis stages (CRN F0-F4) using conventional immunohistochemistry. Finally, functional in vivo validation studies were performed in mice with selective hepatocyte-specific candidate gene deletion to directly assess its requirement for mitochondrial DNA escape and hepatic fibrosis at baseline and under dietary MASH challenge.

**Figure 1.**
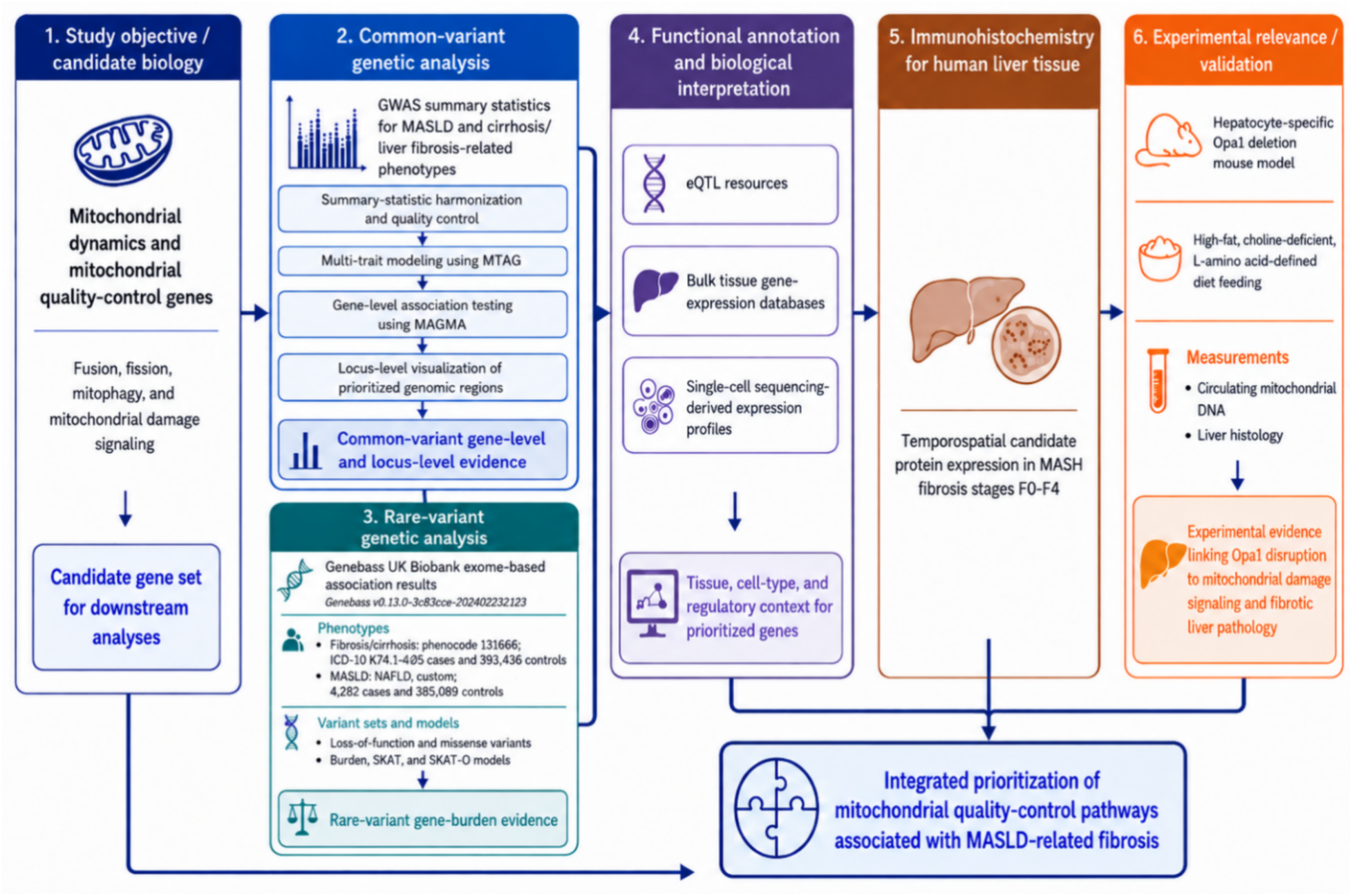
Integrated Workflow to Interrogate Mitochondrial Quality-Control Genetics to MASLD-Associated Fibrosis

### 1.2 Selection of genes for investigation

Candidate target genes (n=35) were selected a priori from two complementary sources: (i) the curated mitochondrial proteome inventory MitoCarta 3.0 (27) and (ii) a targeted original literature review focused on functional studies of mitochondrial fusion, mitochondrial fission, and mitophagy.(28–32) Genes identified from these sources were merged and deduplicated to generate a non-redundant analytical set of mitochondrial dynamics and quality-control genes for downstream gene-based analyses. Genes were then annotated into non-mutually exclusive functional groups, as follows:

**Mitophagy genes** (N = 13): BCL2L13, CALCOCO2, FKBP8, FUNDC1, NIPSNAP1, PARK2, PARK7, PARL, PGAM5, PINK1, TOMM7, USP30, VPS13D.

**Mitochondrial fission genes** (N = 15): ARMC10, DNM1L, FIS1, MFF, MIEF1, MIEF2, MTFP1, MTFR1, MTFR2, MUL1, OMA1, RAB24, SLC25A46, SPIRE1, STX17.

**Mitochondrial fusion genes** (N = 8): ARL2, MFN1, MFN2, MTCH2, OMA1, OPA1, PHB2, PLD6.

Once the previously unrecognized significant association between ***OPA1*** and liver fibrosis in MASLD was identified, we specifically interrogated OPA1-centered mitochondrial quality-control biology by expanding the candidate set to include genes encoding functionally validated OPA1 and OPA1-related pathways direct interactors. The expanded OPA1-related network additionally included the following two categories:

**OPA1 direct-interactor genes** (N = 27): OMA1, YME1L1, PMPCA, PMPCB, MFN1, MFN2, FIS1, SLC25A46, PHB2, PHB1, ATAD3A, HIGD1A, RCC1L, WBSCR16, CHCHD3, MIC19, IMMT, MIC60, SAMM50, PRELID1, TRIAP1, SIRT4, SIRT3, MICU1, TOMM70, PFKFB3, NEDD4L.

**OPA1 pathway genes** (N = 29): DNM1L, MFF, MIEF1, MIEF2, SAMM50, MTX1, MTX2, MICOS10, MICOS13, CHCHD6, APOO, APOOL, TRIAP1, TAMM41, PGS1, CRLS1, TAZ, AGK, PINK1, PRKN, BNIP3, BNIP3L, FUNDC1, CYCS, BAX, BAK1, APAF1, CASP9, CASP3.

SAMM50 and TRIAP1 were present in both the OPA1 direct-interactor and OPA1 pathway categories; each gene was counted once in the non-redundant analytical gene set; a total of 54 unique genes was analyzed.

### 1.3 GWAS summary statistics identification, eligibility, and harmonization

Genome-wide association study (GWAS) summary statistics for MASLD/MASH (including old nomenclature NAFLD- and NASH-related definitions), and liver fibrosis-related phenotypes, including cirrhosis, were identified from the NHGRI-EBI GWAS Catalog (33) and FinnGen releases (34, 35). GWAS datasets were eligible for inclusion if they provided genome-wide summary statistics and sufficient metadata to enable harmonization and multi-trait modeling, including ancestry, sample size, allele coding, and variant-level association information. When multiple GWAS were available for genetically correlated or overlapping phenotypes, we prioritized datasets with the largest effective sample size, well-documented phenotype definitions, and clearly described quality-control procedures.(34, 36–39)

GWAS summary statistics were reformatted into a standardized schema comprising rsID, chromosome, genomic position, effect and non-effect alleles, effect estimate (beta or log-odds ratio), standard error, P value, and sample size. Alleles were harmonized across traits to ensure consistent coding and direction of genetic effects. Variants with missing essential fields were excluded. Strand-ambiguous variants (A/T or C/G) were handled conservatively and removed when allele orientation could not be resolved with confidence.

### 1.4 Multi-trait analysis of GWAS using MTAG

To increase statistical power for locus discovery and downstream gene-based testing of common-variants, GWAS summary statistics for MASLD and cirrhosis/liver fibrosis-related phenotypes (6 GWASs with cumulatively 0.7 million individuals) were harmonized and jointly modeled using Multi-Trait Analysis of GWAS (MTAG).(40) MTAG generates trait-specific association statistics by leveraging shared polygenic architecture across traits while allowing effect estimates to differ by phenotype and accounting for potential sample overlap through the genome-wide covariance structure of the input summary statistics.(40)

Before MTAG analysis, all GWAS summary statistics underwent harmonization and quality control. Analyses were restricted to high-quality autosomal variants with valid effect estimates, standard errors, allele coding, and sample-size metadata. Variants with ambiguous allele coding, unresolved strand orientation, duplicate identifiers, or inconsistent allele frequencies across datasets were excluded. Effect estimates were aligned to a common effect allele to ensure consistent directionality across traits.

We used linkage disequilibrium (LD) score regression to estimate pairwise genetic correlations and cross-trait intercepts, thereby assessing shared genetic architecture, potential sample overlap, and the suitability of each trait combination for MTAG.(41, 42) Trait combinations were retained when they demonstrated interpretable genetic correlation estimates, adequate SNP overlap after harmonization, consistent allele coding, sufficient effective sample size, and no evidence of inflation inconsistent with polygenic signal. Trait pairs were excluded when genetic correlation estimates were unstable, negligible, directionally inconsistent with biological expectations, or when summary-statistic quality was insufficient for reliable multi-trait modeling. These criteria were applied in accordance with established recommendations for MTAG implementation.(40–42) MTAG was then applied to the retained harmonized GWAS summary statistics using its standard modeling framework. Model diagnostics, including trait-specific mean χ² statistics, genetic covariance estimates, and maxFDR, were inspected to evaluate the robustness of MTAG-derived associations.(40) For each phenotype, MTAG produced updated trait-specific association statistics, including effect estimates, standard errors, and P values. These MTAG-derived results were used for downstream locus-level visualization and gene-based association testing. Genome-wide significance was defined as P < 5 × 10⁻⁸ for variant-level analyses.

### 1.5 Gene-based association test of candidate genes

Gene-based association testing of common variants was performed using Multi-marker Analysis of GenoMic Annotation (MAGMA) (43) with trait-specific MTAG-derived summary statistics. SNPs were assigned to genes according to genomic position using a window extending 35 kb upstream and 10 kb downstream of the transcribed region to capture proximal regulatory variation. Gene-level association statistics were calculated using the MAGMA SNP-wise mean model, which aggregates SNP-level evidence while accounting for LD among variants. LD was estimated using an ancestry-matched 1000 Genome Project reference panel when available; otherwise, the closest available reference panel matching the predominant ancestry of the GWAS was used. Analyses were restricted to autosomal genes in the curated mitochondrial dynamics and mitochondrial quality-control gene set.

Rare-variant gene-based associations were evaluated using publicly available Gene-Biobank Association Summary Statistics (Genebass) results from UK Biobank exome-sequencing data, version v0.13.0-43c83cc-202402232123(44). Loss-of-function and missense variant sets were tested for association with MASLD and fibrosis/cirrhosis phenotypes using burden, SKAT, and SKAT-O models. The UK Biobank fibrosis/cirrhosis phenotype was defined using phenocode 131666, corresponding to ICD-10 K74, “fibrosis and cirrhosis of liver,” based on the first occurrence of any mapped three-character ICD-10 K74 code. Any medical event whose recorded event date was earlier than the participant’s date of birth was treated as implausible and removed from the phenotype definition. This phenotype included 1,405 cases and 393,436 controls. The MASLD phenotype was based on the Genebass NAFLD custom definition, derived from a combination of ICD-10, ICD-9, and self-reported codes, and included 4,282 cases and 385,089 controls. Controls for the both phenotypes were quality-control–eligible participants assigned a negative phenotype status—that is, participants who did not meet the corresponding GeneBass case definition; participants with undefined or missing phenotype status were not included as controls. For each candidate gene, we extracted gene-level association results for liver fibrosis and cirrhosis phenotypes across loss-of-function and missense variant masks as implemented in Genebass. Genebass was generated using the SAIGE-GENE mixed-model framework, which supports scalable region- and gene-based testing in large biobank datasets while accounting for case-control imbalance, relatedness, and population structure(45), including complementary aggregation tests, such as SKAT and SKAT-O(46–48). SKAT aggregates qualifying variants within a gene to have heterogeneous directions and magnitudes of effect. SKAT-O adaptively combines burden and SKAT frameworks to improve power across different rare-variant genetic architectures.

For both common- and rare-variant analyses, statistical significance was defined using Bonferroni correction across the number of genes in the curated mitochondrial gene set. Nominal associations were interpreted as supportive evidence and prioritized only when consistent across related phenotypes, variant classes, or complementary gene-based testing frameworks.

### 1.6 Locus-level visualization and regional association plots

Regional association plots were generated using LocusZoom to visualize association signals at loci of interest in their genomic context.(49) For each locus, the plot was centered on the lead variant, defined as the variant with the smallest regional association P value, and displayed surrounding variants, local gene annotations, recombination rates, and LD structure. LD with the lead variant was estimated using an ancestry-matched reference panel when available, or the closest available population reference otherwise. Genomic coordinates were aligned to the genome build of the corresponding GWAS summary statistics. These plots were used to assess whether gene-level associations were supported by localized variant-level signals and to characterize the LD structure surrounding candidate loci.

### 1.7 Functional annotations with GTEx eQTL data

To prioritize candidate effector genes and infer potential regulatory mechanisms, we annotated lead variants and high-LD proxy variants using cis-expression quantitative trait locus (cis-eQTL) data from the Genotype-Tissue Expression (GTEx) Project v8.(23) Proxy variants were selected using ancestry-matched LD estimates where available. For each variant–gene pair, we recorded the tissue context, effect allele, direction of association with gene expression, normalized effect size where available, and corresponding P value or false discovery rate. Particular attention was given to liver and metabolically relevant tissues, while cross-tissue regulatory effects were also considered. These annotations were used to evaluate whether MASLD- or liver fibrosis-associated alleles were linked to altered expression of candidate mitochondrial quality-control genes, thereby helping to distinguish plausible effector genes from nearby genes in LD.

### 1.8 Human liver samples and tissue OPA1 protein expression analysis

De-identified liver biopsy samples from patient with MASLD (age ≥ 18 years; n = 32) were obtained from human sample repository in Tampa General Hospital, Tampa, FL. All patients with MASLD had no other chronic liver disease or significant alcohol consumption (patients with >20 g alcohol daily were excluded from the registry). The study was reviewed and approved by the Institutional Review Board of University of South Florida (Study# 009057). All biopsies were scored according to Kleiner and Brunt et al, and fibrosis was defined according to Clinical Research Network fibrosis staging system as no/minimal (F0–1) or significant (F2–4) (50).

### 1.9 Generation of hepatocyte-specific OPA1 knockout mouse

All mouse experiments were approved by the IACUC of the Beth Israel Deaconess Medical Center (BIDMC, Protocol #003-2021-24). Mice with hepatocyte-selective depletion of OPA1 (OPA1 Hep^KO^) were generated by administering AAV8–thyroxine-binding globulin (TBG)-Cre (AAV.TBG.PI.Cre.rBG, Addgene #107787) via an intraperitoneal (i.p.) injection at 1.5 × 10^11^ genome copies into Opa1^fl/fl^ mice on a C57BL/6-129/SvEv mixed background(51). Equivalent dose of AAV8-TBG-GFP (AAV.TBG.PI.eGFP.WPRE.bGH, Addgene #105535) was administered as control. Upon arrival to SPF animal facility at BIDMC, OPA1^fl/fl^ were backcrossed to C57Bl6J (Jackson Labs) for additional 2 generations and intercrossed to obtain homozygous breeders to establish a colony. Male mice were used in all experiments starting from age 8-10 weeks.

### 1.10 Dietary murine model of progressive fibrotic MASH

High-fat choline-deficient, amino acid defined diet (HF-CDAA, A06071302, Research Diets) was fed ad libitum to induce aggressive MASH-like steatohepatitis, characterized by ductular reaction, progressive hepatic fibrosis eventually resulting in cirrhosis and HCC, as previously described in detail(52). Regular rodent chow (Purina D5008) was fed to age- and sex-matched to non-MASH controls. HF-CDAA or control chow diet was initiated in male OPA1^fl/fl^ five days after AAV8-mediated Cre recombination and continued for 8 weeks, when HF-CDAA animals develop advanced steatohepatitis and fibrosis, before euthanasia and tissue analysis.

### 1.11 Histology and immunohistochemistry

Connective tissue stain (Sirius Red), Hematoxylin-eosin (HE) and IHC for alpha-SMA were performed in formalin-fixed paraffin-embedded murine liver sections, as described previously(52). Immunohistochemistry for OPA1 was performed in human FFPE sections using conventional DAB immunohistochemistry protocol after Tris-EDTA Buffer (pH 9.0) antigen retrieval. Primary rabbit anti-OPA1 antibody were purchased from Sigma-Aldrich (HPA036926, 1:100 dilution).

### 1.12 Digital droplet PCR (ddPCR) mtDNA assay

Cell-free, circulating DNA was isolated from 50ul of EDTA plasma samples after additional centrifugation at 13,000g for 10 minutes at 4C using the QIAamp DNA Blood Kit (Qiagen, Valencia, CA). The absolute concentration of the mtDNA template (copy numbers/μL of plasma) was quantified using the TaqMan principle and previously published primer/probes labelled with hexachlorofluorescein- or 6-carboxy-fluorescein (12) on the QX200 ddPCR System (Bio-Rad) via simultaneous amplification of sequences within 12S region of the mtDNA genome and b2-microglobulin within nuclear DNA.

### 1.13 Software, reproducibility, and reporting

Analyses were conducted using published, validated, and publicly available software. LDSC regression (LD SCore v1.0.1; github.com/bulik/ldsc), MTAG (github.com/JonJala/mtag) and MAGMA (cncr.nl/research/magma/) were run using their recommended input formats and default settings unless otherwise specified. Downstream analyses and visualization were performed in R (version 4.4.2) or Python (version 3.13.13) using standard scientific computing libraries, including LDlinkR (github.com/CBIIT/LDlinkR), locuszoomr (github.com/myles-lewis/locuszoomr), EnsDb.Hsapiens.v75 (www.bioconductor.org/packages/release/data/annotation/html/EnsDb.Hsapiens.v75.html), and pandas (pandas.pydata.org). All parameter choices (gene windows, correction thresholds, and reference panels) were reported to enable reproducibility.

## Results

### 1.1 Multi-trait GWAS increases discovery power and recapitulates established MASLD genetic architecture

MTAG integrated six large-scale GWAS datasets with significant cross-trait genetic correlation (**Figure 1A**; **Supplementary Table 1**; P < 0.05). LDSC metrics supported adequate control of confounding, with intercepts near 1.0 (0.998–1.027) and attenuation ratios largely <0.20–0.30, indicating that observed signal inflation was primarily due to polygenicity rather than bias (**Supplementary Table 1**).

MTAG yielded equivalent sample size of 54,194 to 1.8 million and tested ∼550 million variants across the included GWASs. Among these GWASs, MTAG-adjusted GWAS summary statistics for liver cirrhosis (GCST90319878 with mixed ethnicity), jointly analyzed with MASLD summary statistics has the largest equivalent sample size of 1.8 million, and therefore, was used in subsequent analysis. This MTAG-adjusted GWAS identified 1,277 genome-wide significant variants mapping to 25 independent loci (**Figure 2C**), reflecting increased discovery power compared to single-trait GWAS.

**Figure 2.**
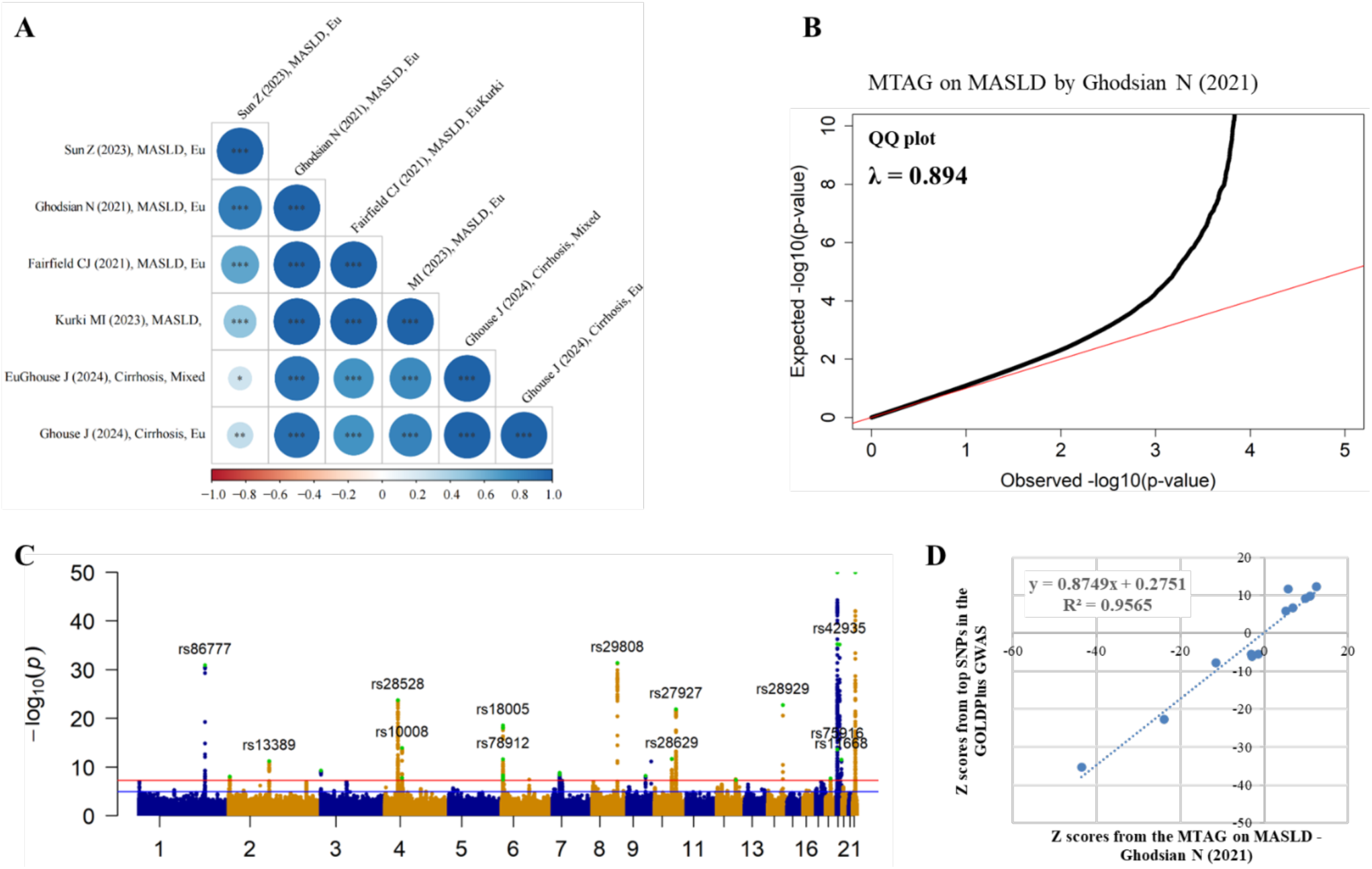
Multi-trait analysis of GWASs increased discovery power and recapitulates established MASLD genetic architecture. A. The six selected GWASs were genetically correlated and were included in the MTAG. B. The QQ plot showed marked deviation from the null consistent with polygenic architecture, while the genomic inflation metric (λ ≈ 0.894). C. Manhattan plot of the MTAG on the GWAS on MASLD. D. MTAG results showed strong correlation with the known MASLD-associated variants reported by the GOLDPlus meta-analysis.

Quality-control analyses supported robust signal enrichment. The QQ plot showed deviation consistent with polygenic architecture (**Figure 2B**), and the genomic inflation factor (λ = 0.894) indicated no systematic inflation. Per-variant effect estimates were highly concordant with previously reported MASLD/NAFLD loci from the GOLDPlus meta-analysis(53) (R² > 0.95; **Figure 2D**), confirming that the multi-trait framework recapitulates established MASLD genetic architecture while enabling additional locus discovery.

### 1.2 Gene-based association analysis implicates OPA1 in liver fibrosis-related traits

Gene-level association analysis of common variants across 35 genes involved in mitochondrial dynamics and quality control identified enrichment of liver fibrosis-associated signals at the *OPA1* locus. In the combined MTAG-MAGMA analysis for liver fibrosis (accession GCST90319878, mixed-ancestry), *OPA1*—which encodes a key mediator of inner mitochondrial membrane fusion and cristae remodeling—showed the strongest gene-level association, surpassing the Bonferroni-corrected significance threshold (P = 1.1 × 10⁻³, corrected threshold ≈1.3 × 10⁻³). Conditional analysis excluding the lead single-nucleotide polymorphisms within the *OPA1* locus retained a significant association with the liver fibrotic phenotype (P < 0.05), suggesting the presence of multiple independent signals. Locus visualization revealed a cluster of association signals that co-localize with the local LD structure (**Figure 3**).

**Figure 3.**
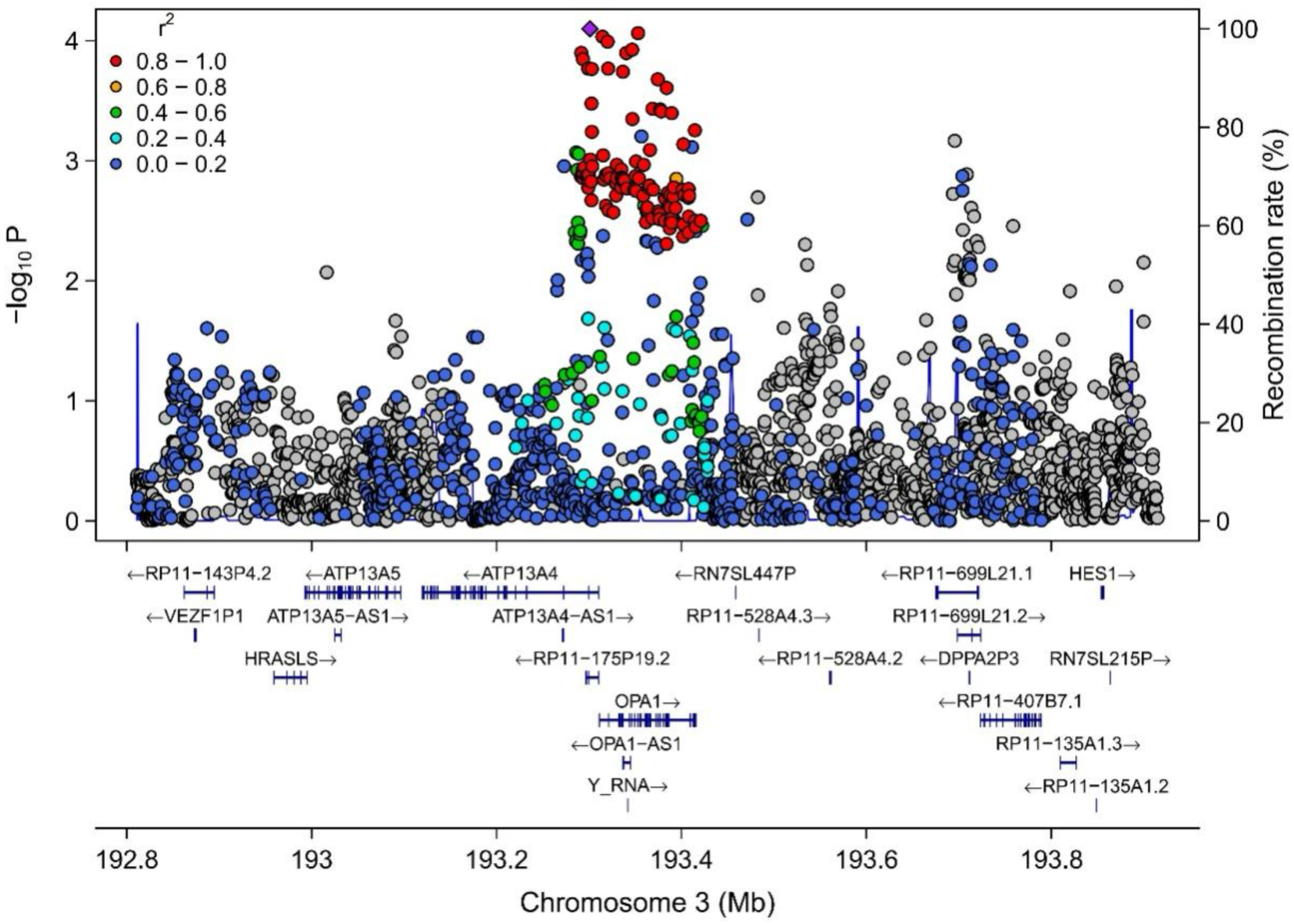
*OPA1* locus regional association plot with local LD, recombination rate, and gene annotations. Regional association signals surrounding *OPA1* on chromosome 3 are plotted as −log₁₀(*P*) against genomic position. Variants are colored according to linkage disequilibrium (LD; *r*²) with the index variant, as indicated by the legend. The local recombination rate is shown on the right axis, and gene locations and transcriptional orientations are displayed below the association plot.

Several other genes demonstrated nominal gene-level associations in the MTAG analysis, including the mitochondrial fusion-related gene *MTCH2* (P = 0.033), the fission-related gene *MIEF1* (P = 0.0047), and the mitophagy-related gene *CALCOCO2* (P = 0.05) (**Figure S1**).

To further evaluate the contribution of *OPA1* to individual predisposition to liver fibrosis, we examined rare coding variants using next-generation sequencing data from the UK Biobank. Gene-based collapsing analysis of loss-of-function variants in *OPA1* revealed a borderline nominally significant association with liver fibrosis and cirrhosis by SKAT (P=0.05), but not with metabolic dysfunction-associated steatotic liver disease (MASLD; P≥0.33), supporting specific role in liver fibrogenesis (**Table 2**).

**Table 1.**
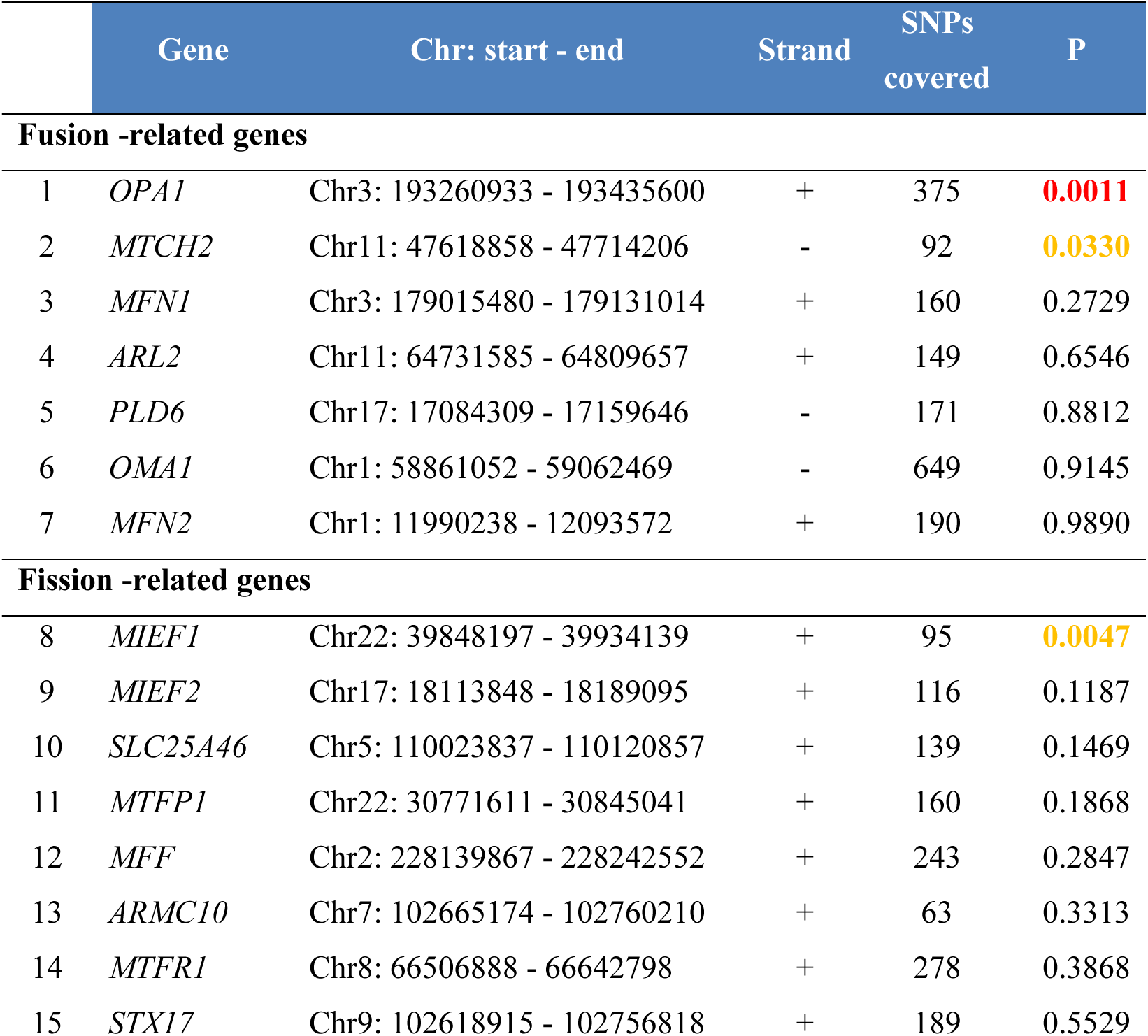
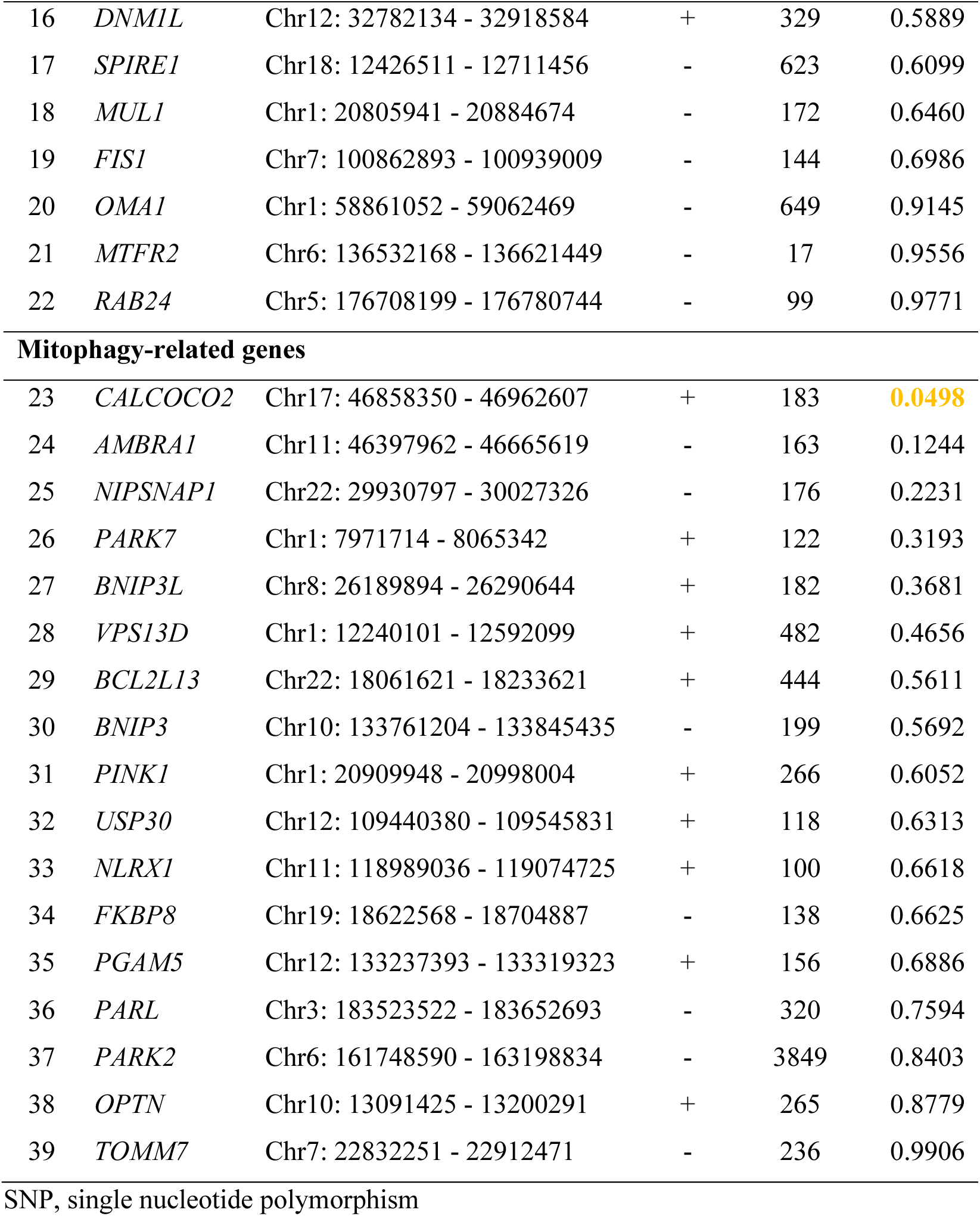
MTAG-based gene-level association of common variants in mitochondrial dynamics and quality-control genes with liver fibrosis.

**Table 2.** Rare-variant burden analysis of *OPA1* for liver fibrosis, cirrhosis, and MASLD in UK Biobank sequencing data.

| Phenotype | Sample size<br>(cases vs. control) | Type of<br>variant | Test | Beta | P value |
| --- | --- | --- | --- | --- | --- |
| Fibrosis and<br>cirrhosis of<br>liver | 1,405 vs. 393,436 | LoF | Burden | 0.0626 | 0.308 |
|  |  | LoF | SKAT | 0.0626 | **0.050** |
|  |  | LoF | SKAT-O | 0.0626 | 0.087 |
|  |  | Missense | Burden | 0.0051 | 0.503 |
|  |  | Missense | SKAT | 0.0051 | 0.780 |
|  |  | Missense | SKAT-O | 0.0051 | 0.723 |
| MASLD | 4,282 vs. 385,089 | LoF | Burden | -0.0406 | 0.318 |
|  |  | LoF | SKAT | -0.0406 | 1.000 |
|  |  | LoF | SKAT-O | -0.0406 | 0.522 |
|  |  | Missense | Burden | 0.0044 | 0.325 |
|  |  | Missense | SKAT | 0.0044 | 0.339 |
|  |  | Missense | SKAT-O | 0.0044 | 0.514 |

### 1.3 Gene-based association of OPA1-interacting and pathway genes with liver fibrosis

To further explore the contribution of *OPA1* and its network to liver fibrosis, we examined common variants in 41 additional genes encoding known *OPA1* interactors and pathway components, using multi-trait GWAS summary statistics (MTAG; accession GCST90319878, mixed-ancestry). *SAMM50* showed a robust association with liver fibrosis risk (P = 1.96 × 10⁻¹⁶), and both *MIEF1* (P = 0.0047) and *CHCHD3* (P = 0.050) reached borderline nominal significance (**Table 3**).

**Table 3.**
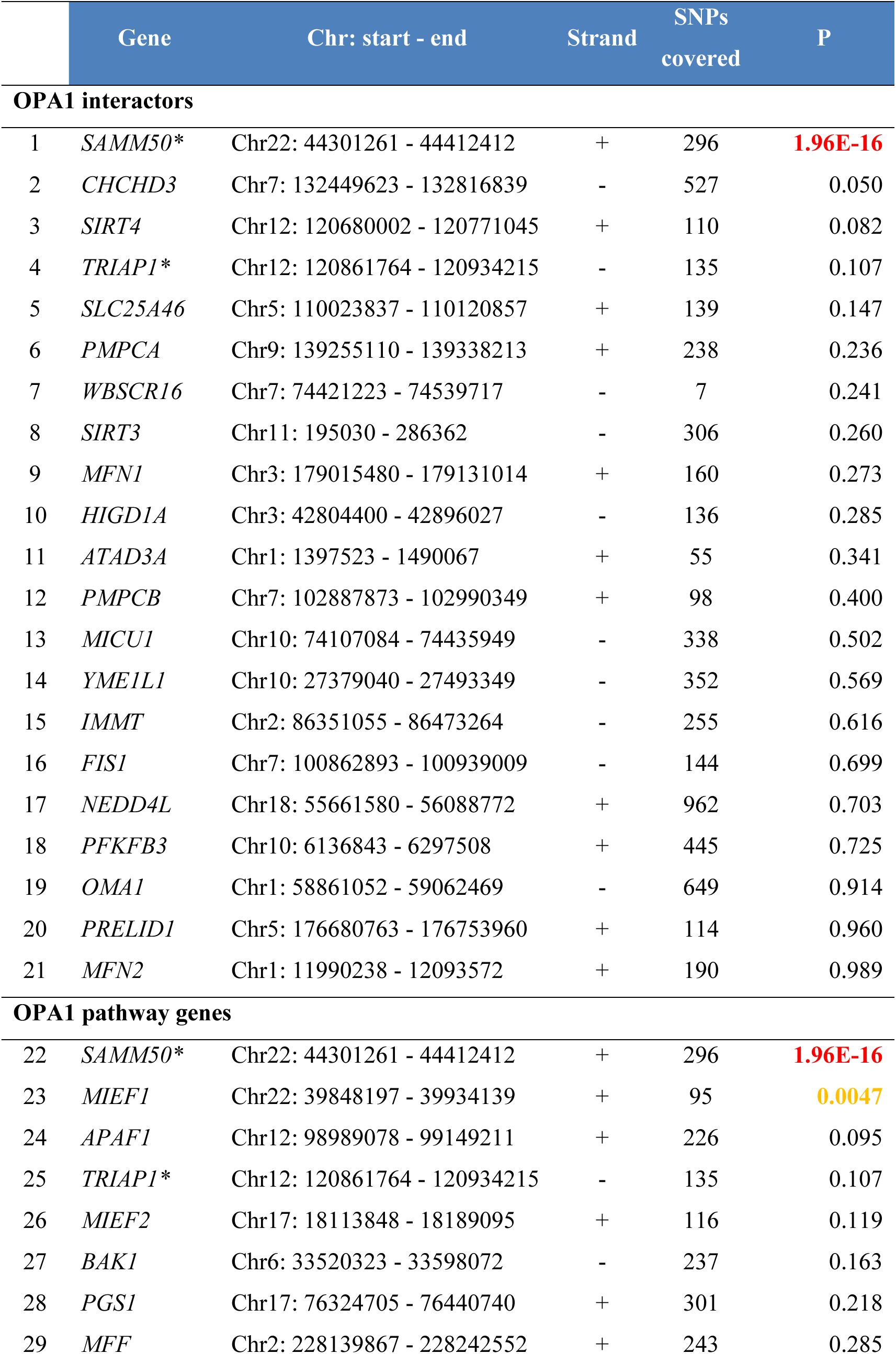
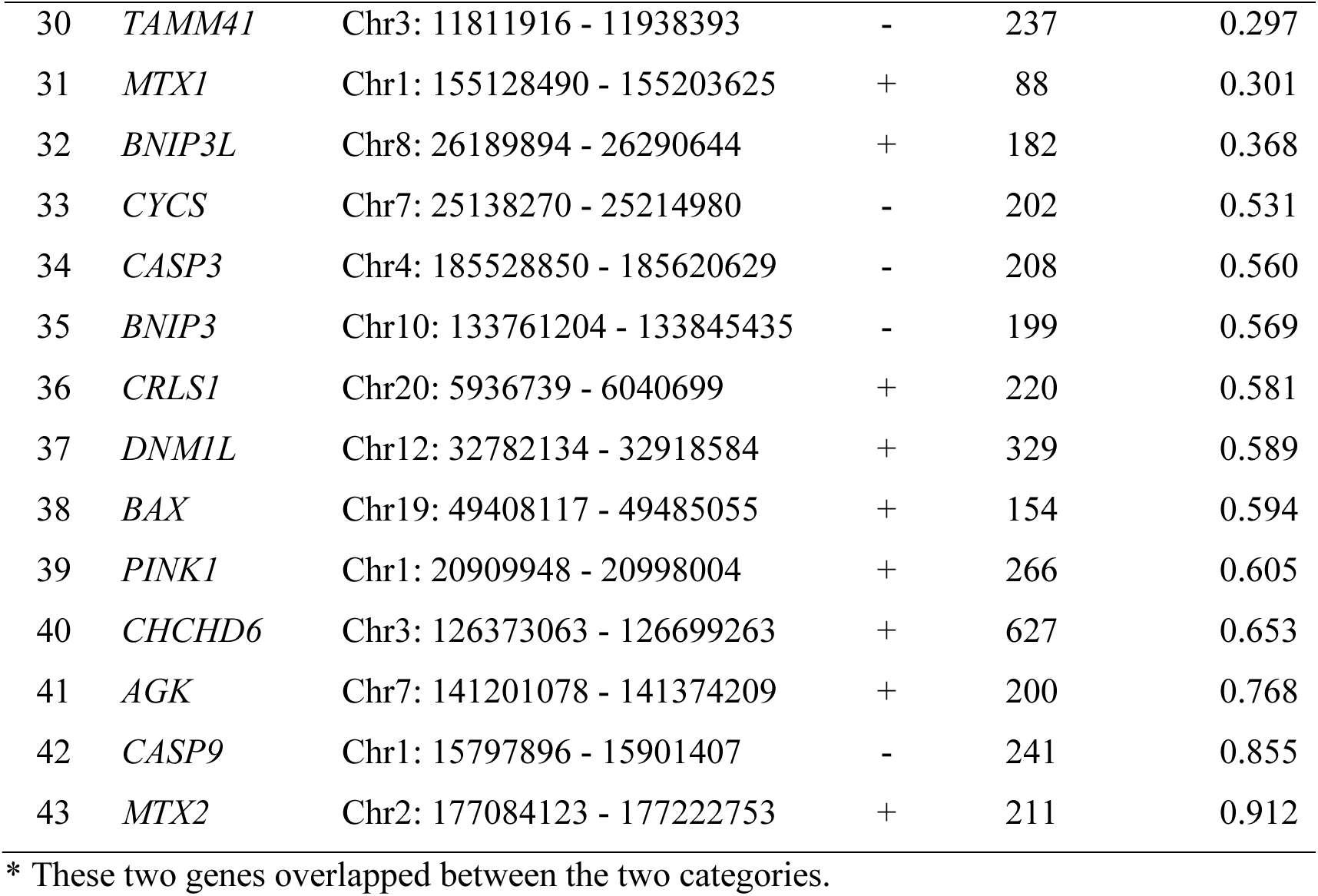
Common-variant gene-based test of *OPA1* interactors and pathway genes using MAGMA.

In gene-based burden tests of rare variants, we found nominally significant enrichment of loss-of-function or missense variants in seven *OPA1*-related genes for liver fibrosis and cirrhosis (P ≤ 0.047; **Table S2**) and in a distinct set of seven genes for MASLD (P ≤ 0.045; **Table S3**). Among these, *CRLS1* for fibrosis/cirrhosis and *BAK1* and *CYCS* for MASLD were nominally significant across all three burden test models. No rare-variant association, however, survived Bonferroni correction for multiple testing. Collectively, these findings indicate that proteins interacting with *OPA1* and components of *OPA1*-related pathways may modulate susceptibility to liver fibrosis. The nominal *OPA1* and related gene rare-variant burden association likely reflects limited statistical power due to the scarcity of qualifying variants and low carrier counts. The stronger signal for fibrosis/cirrhosis than for MASLD further suggests that *OPA1*-related mitochondrial dysfunction may preferentially contribute to fibrotic progression rather than disease initiation. Interestingly, *SAMM50* showed the strongest gene-based association ( *P* = 1.96 × 10^−16^); however, *SAMM50* resides within the established *PNPLA3–SAMM50* susceptibility region, where extensive regional association and linkage disequilibrium complicate gene-level attribution. We therefore interpreted this signal at the locus level and prioritized OPA1 for subsequent investigation based on convergent evidence from independent genetic and tissue-level analyses.

### 1.4 MTAG-identified OPA1 locus variants show eQTL evidence for regulation of OPA1 expression

To assess the biological relevance of the variants in the *OPA1* locus for regulation of gene expression, we evaluated eQTL evidence. Multiple significant MTAG variants (P≤9.26×10^-5^) within or near *OPA1* were significantly associated with *OPA1* eQTL variants across different tissues which include directly relevant ones, such as *cultured fibroblast* and *whole blood* (**Table 4 and Table S4**). Specifically, the most significant signals included variants located in *OPA1* intron 1 and upstream regulatory regions, with corresponding high-LD eQTL proxies (R² > 0.7) demonstrating significant expression effects (P ≤ 7.0×10^-5^). Notably, SNPs rs9856156 and rs4597663 are ranked as 1f in the RegulomeDB (version 2.2) database, suggesting strong experimental evidence supporting their gene expression regulatory effects.

**Table 4.**
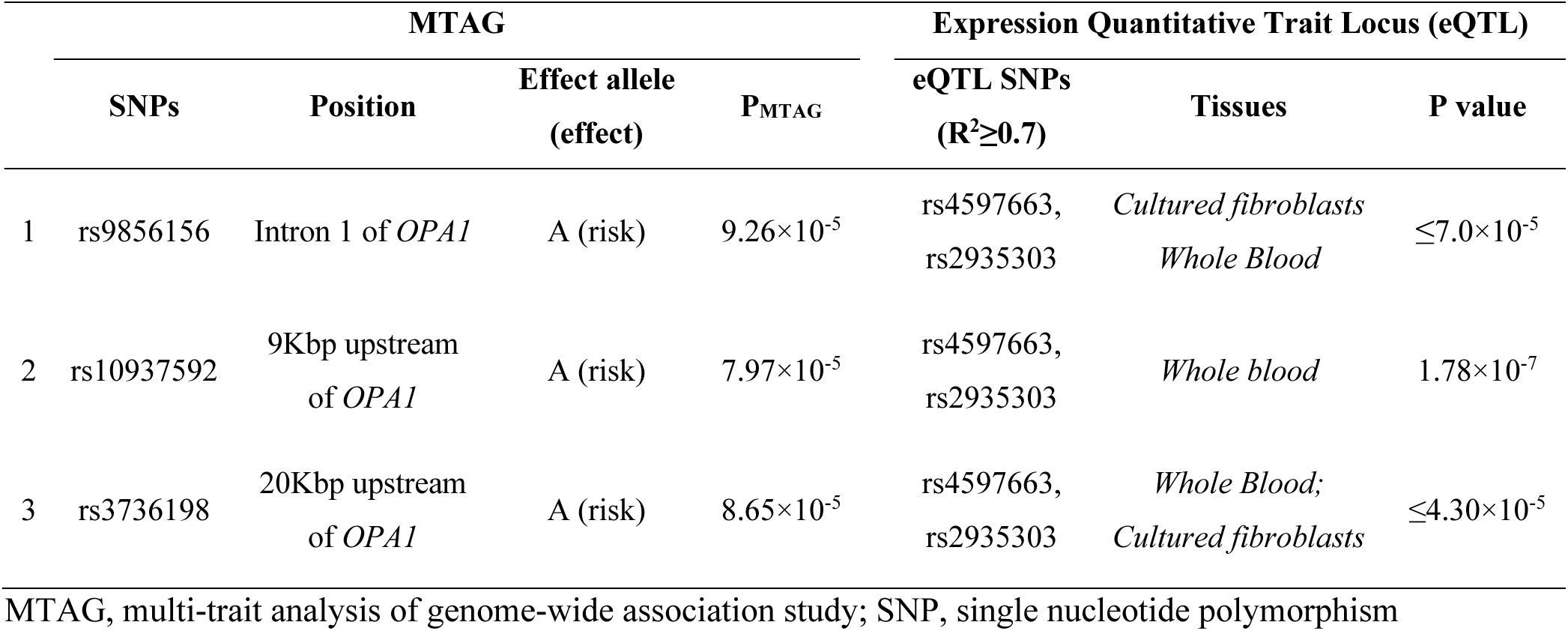
Significant MTAG variants in or near *OPA1* have regulatory effects on OPA1 expression.

### 1.5 OPA1 shows liver-relevant expression and disease-associated dysregulation in MASLD and cirrhosis

OPA1 is constitutively expressed in human liver at both transcript and protein levels: Human Protein Atlas data show moderate hepatic protein staining, and *OPA1* RNA ranks fifth among the surveyed tissues (**Figure S2**).(24, 25) Published human transcriptomic studies further indicate higher hepatic *OPA1* mRNA in MASLD, with positive associations with NAFLD Activity Score in two independent cohorts (GSE135251 and GSE207310).(54) In contrast, protein-level findings are not directionally uniform. Western blotting in a 130-patient bariatric cohort showed reduced hepatic OPA1 protein from NAFL onset through NASH progression,(55) whereas an independent DIA-MS liver proteome cohort found no significant change in total OPA1 abundance in NASH versus controls.(56) These studies therefore support OPA1 dysregulation in MASLD but demonstrate that transcript abundance and total protein do not necessarily change in parallel (**Table S5**). Evidence in advanced fibrosis/cirrhosis is more limited and similarly argues against a simple monotonic expression model. In the same DIA-MS cohort, total hepatic OPA1 abundance was not significantly altered in cirrhosis versus controls,(56) while older HCC-centered tissue data suggested that OPA1 reduction may already occur in cirrhotic liver but did not provide a definitive cirrhosis-versus-normal estimate.(57) More recent mechanistic work showed that fibrosis can alter *OPA1* splicing and deplete functional long-form OPA1 without requiring a major change in total OPA1 abundance.(58) Collectively, the literature is most consistent with disease-associated OPA1 dysregulation—encompassing RNA abundance, total protein, and isoform/processing changes—rather than a uniform increase or decrease in OPA1 expression across MASLD progression and cirrhosis.

### 1.6 Histological analysis in human MASH reveals temporospatial change in OPA1 protein expression parallels fibrosis progression

We have performed a survey of human MASH liver tissue expression of OPA1 at a protein level via immunohistochemistry in 32 archival tissue specimens at various fibrosis stages (F0, n=6; F1, n=7; F2, n=6; F3, n=5; F4, n=8). In non-fibrotic F0/1 livers, moderate OPA1 immunopositivity was observed in hepatocytes, which increased progressively from F2 to F4 fibrosis stages (**Figure 4A**). Interestingly, markedly increased OPA1 signal in advanced fibrosis stages was observed largely within liver epithelium – primarily hepatocyte, but also cholangiocyte-like cells and ductular structures. Faint OPA1 staining in non-fibrotic hepatocytes became overall more intense in advanced fibrosis, and demonstrated significant cell-to-cell variation in expression level. Cholangiocytes within bile ducts in non-fibrotic MASH were negative, whereas in advanced fibrosis stages proliferating duct-like structures (reactive cholangiocytes/hepatic progenitors, “ductular reaction”) were positive (**Figure 4B,C**).

**Figure 4.**
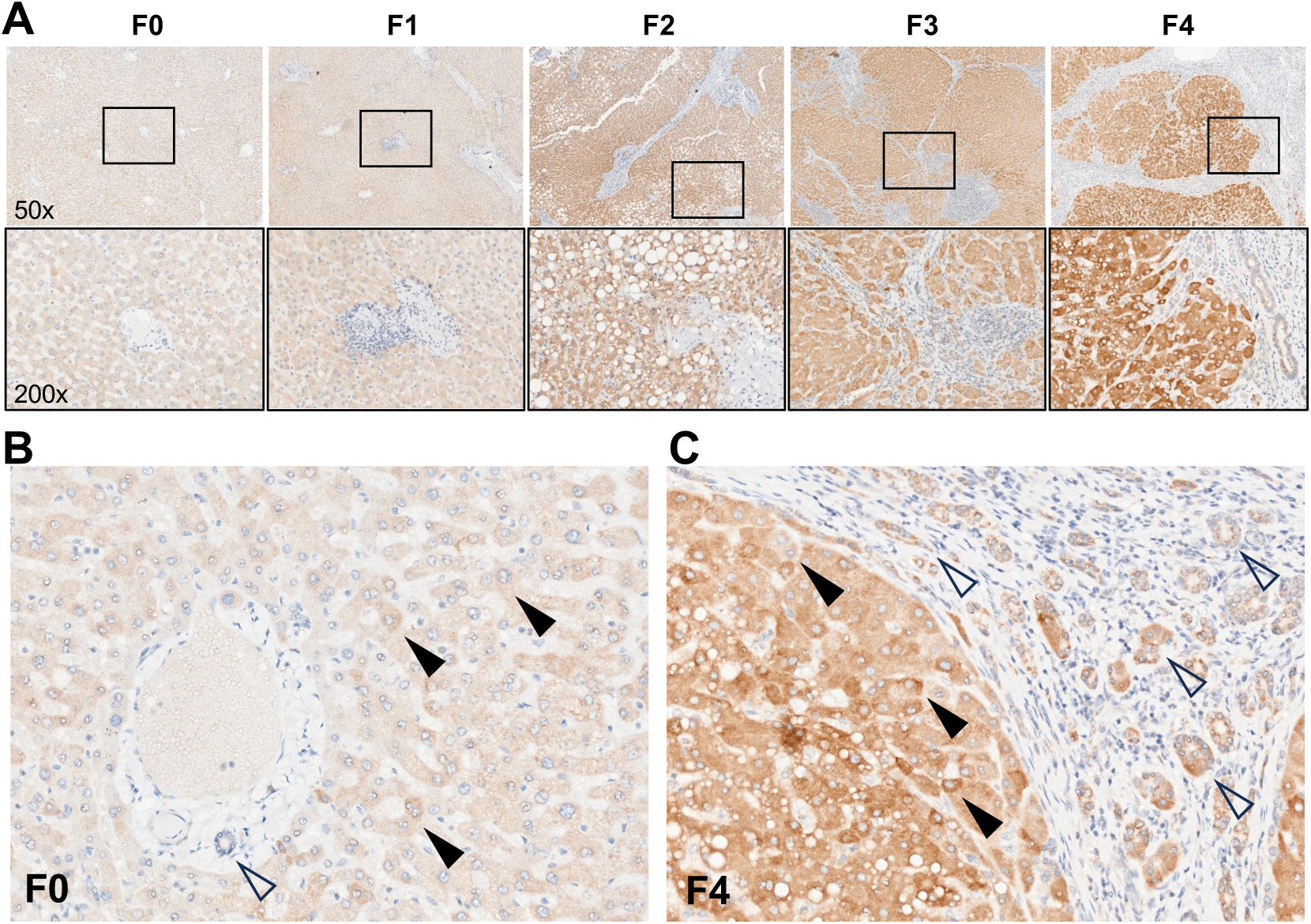
Changes in OPA1 protein expression in relation to fibrosis stage in human MASH. Immunohistochemistry for OPA1 was performed in 32 human liver FFPE sections as described in M&M. (A) Representative staining of liver tissue from MASH patients with progressive fibrosis stages (CRN F0-F4). Upper panel are low magnification images (x50), and lower panel are blow-up magnification (x200). (B-C) Representative high-magnification images of non-fibrotic MASH (B, F0) and end-stage cirrhosis (C, F4) demonstrate marked increase in OPA1 expression in hepatocytes and duct-like structures within fibrotic septa in MASH cirrhosis. Hepatocytes show variable immunopositivity for OPA1 (closed arrows, and bile ducts and ductular proliferations (pseudo-ducts) are marked by open arrows.

### 1.7 Loss of OPA1 in hepatocytes promotes mito-DAMPs escape and fibrogenic activation of hepatic stellate cells in vivo

Since OPA1 expression changes are predominantly observed in hepatocytes in human MASH, we have performed a hepatocyte-specific OPA1 gene knockout in hepatocyte using floxed OPA mouse injected with TBG.AAV-Cre. In healthy mice fed regular chow, OPA1 deletion alone was sufficient to induce hepatic stellate cell activation, as assessed via alpha-SMA immunohistochemistry and resulted in moderate, but readily detectable zone 3 fibrosis with frequent central-central bridging and thin fibrotic septa formation (**Figure 5A**). We also observed previously reported weight loss(59, 60), and serum liver injury markers ALT and AST were increased without hepatomegaly in OPA1 Hep^KO^ (Figure S3A-D). There was a trend to elevated average circulating mtDNA levels as measured by ddPCR in plasma (15.76±2.13 copies/ul compared to 10.8±3.96, not significant, Figure S3C).

**Figure 5.**
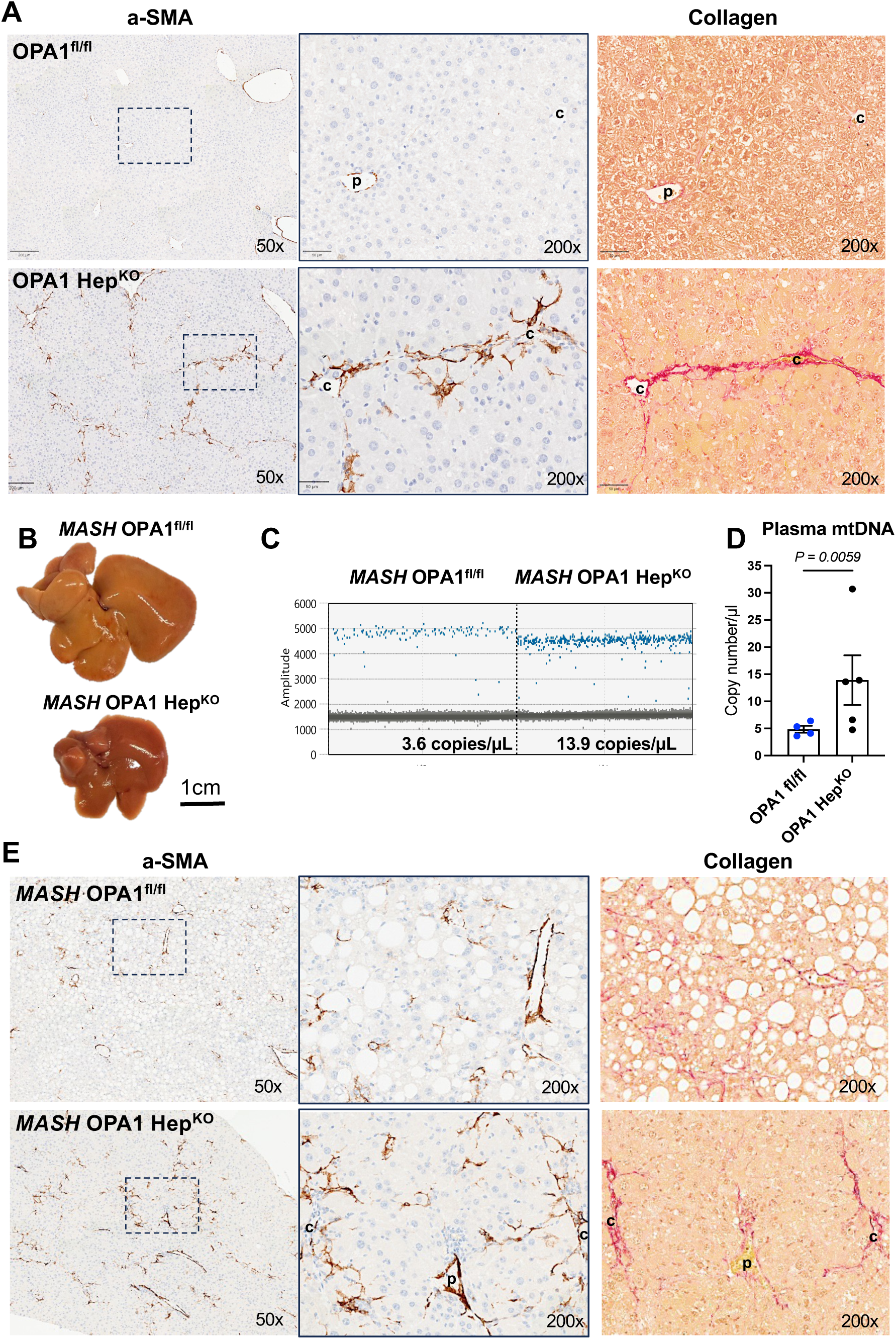
Hepatocyte-specific deletion of OPA1 triggers spontaneous stellate cell activation and hepatic fibrosis, and increase circulating mtDNA levels in mice with experimental MASH. Hepatocyte-specific deletion of OPA1(OPA1 Hep^KO^) was achieved by AAV8-TBG-Cre injection into floxed OPA1 mice. Equivalent dose of AAV8-EGFP was used in OPA1-sufficient control group (OPA1^fl/fl^). Mice were fed regular chow (A, n=5) or HF-CDAA diet (MASH, B-E, n=4-5) for 8 weeks before analysis. (A) Representative immunohistochemistry images of hepatic stellate cell activation marker alpha-SMA at low magnification (x50, left panel) and blow-out high magnification of indicated area (x200, middle panel). Serial section was stained for collagen (picrosirius red, right panel). (B) Representative gross macroscopic images of livers in OPA1 Hep^KO^ and control mice fed MASH diet for 8 weeks. (C) Representative digital droplet PCR amplitude plots for 12S mtDNA fragment and (D) quantification of circulating mtDNA levels in plasma of OPA1-deficient mice and respective controls. (F) Representative immunohistochemistry images of hepatic stellate cell activation marker alpha-SMA at low magnification (x50, left panel) and blow-out high magnification of indicated area (x200, middle panel). Serial section was stained for collagen (picrosirius red, right panel). “p” indicates portal vein; “c” indicates central vein.

Next, we performed MASH diet challenge studies after hepatocyte-specific OPA1 deletion followed by 8 weeks of HF-CDAA diet feeding. Despite previously reported (59, 60) weight loss, reduced hepatomegaly and remarkable protection from liver steatosis (**Fig. 5B**, Figure S3E-G) induced by OPA1 loss, hepatocyte-specific OPA1-/- mice demonstrated markedly increased circulating extracellular mtDNA levels in plasma (**Fig. 5C,D**) and exaggerated fibrogenic activation of hepatic stellate cells (**Fig. 5E**). Interestingly, serum ALT decreased in OPA1 Hep^KO^, consistent with protection from steatosis-induced lipotoxicity, but AST was at the same time elevated. Connective tissue staining revealed that OPA1 deficiency resulted in a major shift in scarring pattern from diffuse, pericellular “chicken-wire” fibrosis to compact, zone 3 fibrosis resembling septa-like lesions observed in OPA1 Hep^KO^ at baseline (**Fig. 5E**). At both baseline and under MASH challenge, total bilirubin levels were not affected by hepatocyte loss of OPA1.

## Discussion

Using large-scale MAGMA-MTAG analysis of human genomic datasets focusing on mitochondrial genes, we identified a previously unrecognized significant association between ***OPA1*** and liver fibrosis in MASLD. Complementary analyses of single-cell expression datasets and immunohistochemistry in human liver tissue revealed epithelial OPA1 dysregulation on both mRNA and protein level in MASLD linked with fibrosis progression. Subsequent functional interrogation in mice with hepatocyte-specific OPA1 loss showed striking spontaneous HSC activation and fibrosis in zone 3. Under dietary MASH challenge, hepatocyte OPA1 deficiency resulted in increased circulating mtDNA levels and exaggerated fibrogenic responses despite remarkable protection from liver steatosis and hepatomegaly, suggesting OPA1 plays an indispensable role in preventing pro-fibrogenic mito-DAMP (mtDNA) release and hepatic fibrosis. Together, these findings position OPA1 as a central genetic and biological anchor linking mitochondrial quality-control pathways to mito-DAMPs escape and liver fibrosis in MASLD. Further investigation of this novel pathway may help identify specific pharmacological targets and develop innovative mitochondria-targeted therapeutic strategies for progressive MASLD.

The importance of OPA1 in human disease has been recently firmly established outside the liver: OPA1 mutations cause autosomal dominant optic atrophy or more severe multisystem disease (dominant optic atrophy plus) characterized by deafness, neuropathy, myopathy, and other neurological manifestations(61, 62) but hepatic involvement has not been described in these foundational OPA1 cohorts. OPA1 gene encodes a highly conserved mitochondrial inner-membrane dynamin-related GTPase that regulates inner-membrane fusion, cristae architecture, respiratory function, mtDNA maintenance, and resistance to apoptosis - processes that may determine whether stressed hepatocytes preserve mitochondrial homeostasis or release immunostimulatory mito-DAMPs (63–66). However, studies on OPA1 significance in the liver produced somewhat complex and seemingly contradictory results. Under high-fat feeding challenge, OPA1Hep^KO^ were reported by Da Dalt et al. to be paradoxically protected from steatosis after 20 weeks of feeding (67). Lee et al. found that loss of OPA1 in hepatocytes was dispensable for mitochondrial respiration despite altered cristae morphology in healthy adult mouse liver, and protected against APAP-induced hepatotoxicity (59). One would predict hepatocyte OPA1 deficiency should worsen steatosis, because OPA1 is required for normal oxidative phosphorylation, fatty acid oxidation, and mitochondrial integrity. Surprisingly, these studies and our own results (Fig. 5) show that hepatocyte-specific OPA1 deletion reduces hepatic triglyceride accumulation during high-fat feeding, despite causing mitochondrial dysfunction. This complex phenotype complicates assessment of downstream fibrotic responses and appears to result from systemic metabolic adaptation, rather than healthier hepatocyte mitochondria. The link between OPA1 deficiency, mitochondrial stress signaling, and hepatokine hormone FGF21 was first established by Pereira et al. in skeletal muscle(68) and subsequently in brown adipose tissue(69), by demonstrating that inducible OPA1 loss causes mitochondrial dysfunction and activates an integrated stress response (ISR), robust FGF21 production and systemic protection from diet-induced obesity and insulin resistance. Lee et al. established this pathway directly in the liver, showing that hepatocyte OPA1 deletion induces eIF2α phosphorylation and hepatic FGF21 expression, increases circulating FGF21, and triggers a broader mitohormetic program associated with mitochondrial biogenesis and systemic metabolic adaptation(59). Perhaps due to such striking protection from steatosis, and absence of overt hepatocyte injury and death exaggerated fibrotic responses in OPA1 deficient mice we observe in our study were overlooked in earlier studies.

Large scale genetics analysis reported here identify OPA1 as a genetic node linked to fibrosis in MASH, and may explain how mitochondrial dysfunction in MASH leads to mito-DAMPs escaping injured hepatocytes and triggering fibrotic response(12). OPA1-mediated disruption of mitochondrial cristae can promote mtDNA escape and inflammatory sensing through cGAS–STING-related pathways,(70) providing a mechanistic link between impaired mitochondrial dynamics, innate immune activation, and fibrogenesis. In this context, the gene-based association of OPA1 with fibrosis-related traits, together with signals among OPA1-interacting and mitochondrial dynamics genes, supports the concept that inherited variation in mitochondrial fusion and stress-response pathways may alter susceptibility to progressive fibrosis. The eQTL evidence further suggests that risk-associated variants may influence OPA1 expression, while liver single-cell data showing liver-relevant OPA1 expression and reduced OPA1 signal in MASLD provide tissue-level plausibility. Biologically, altered OPA1 function could promote fibrosis by increasing hepatocyte mitochondrial injury, mtDNA instability, ROS generation, cytochrome c release, and mito-DAMP exposure, thereby amplifying macrophage activation and stellate-cell fibrogenic programs.(12, 18, 19, 70) Mitochondrial dynamics may also act directly within hepatic stellate cells, where mitochondrial fission and oxidative phosphorylation have been linked to stellate-cell activation and experimental fibrosis.(71) Importantly, liver-specific OPA1 perturbation can produce context-dependent adaptive responses in experimental models,(59, 67) indicating that OPA1 is unlikely to act as a uniformly protective or deleterious factor. Rather, our data support OPA1 as a fibrosis-modifying mitochondrial node whose effects may depend on cell type, metabolic stress, and disease stage. Together, these findings position OPA1-dependent mitochondrial architecture as a plausible genetic and biological bridge linking mitochondrial injury, mito-DAMP signaling, immune–stromal crosstalk, and fibrotic progression in MASLD. OPA1 and its related mitochondrial quality-control genes appear most relevant to *fibrosis progression*, particularly the transition from steatotic liver injury to clinically significant or advanced fibrosis, rather than to isolated steatosis or very early scar formation. This interpretation is supported by multiple lines of evidence of OPA1 biology and the phenotype structure of our analyses. OPA1 maintains mitochondrial inner-membrane fusion, cristae integrity, respiratory efficiency, mtDNA stability, and resistance to apoptosis-related mitochondrial stress.(59, 63, 64, 66, 72) Therefore, altered OPA1 function is most likely to become pathogenic when chronic metabolic stress overwhelms mitochondrial adaptation, promoting mito-DAMP release and hepatic stellate cell activation. Consistent with this model, the MTAG-derived signal and rare-variant burden associations were observed in liver fibrosis/cirrhosis-related phenotypes, which more closely reflect advanced, progressive fibrotic disease rather than early F0–F1 fibrosis stages. Perhaps most direct evidence of OPA1 as a key molecular link between mitochondrial dysfunction and fibrosis comes from our striking observation that OPA1 loss in hepatocytes alone cause persistent activation of stellate cells and bridging fibrosis clearly localizing to zone 3 in liver lobule (**Figure 5**); such scarring pattern is characteristic since fibrosis in adult MASH begins as perisinusoidal/pericellular fibrosis in the centrilobular (zone 3) compartment and subsequently extends to portal areas and bridging septa.(50: Brunt, 1999 #2498) An interesting possibility in this model is a OPA1-deficiency dependent release of mito-DAMPs without hepatocyte death, generally assumed to be a requirement(12). Despite the fact that OPA1 loss caused known increase in hepatocyte cell injury marker ALT at baseline (Figure S3C), Lee et al previously showed via liver proteomics an approximately 8-fold increase in hepatic ALT protein in OPA1 Hep^KO^ in the absence of histological injury and argued that the elevated circulating transaminases may partly reflect altered enzyme abundance caused by OPA1 deletion rather than hepatocyte death.(59) Furthermore, phenotype in OPA1 Hep^KO^ after 8 week MASH diet challenge corroborates even stronger the pathogenetic sequence OPA1 loss → mitochondrial disfunction → mito-DAMPs release → HSC activation → fibrosis, because these manifest despite remarkable protection from main disease drivers: steatosis and hepatocyte injury induced by HF-CDAA feeding, as evidenced by robust decrease in (cytosolic) ALT (Figure S3G). Intriguingly, here (mostly mitochondrial) AST demonstrates divergent pattern of elevation, most likely due to molecular dysfunction secondary to OPA1 loss (Figure S3G). It is tempting to speculate that this may reflect the unrecognized mechanistic underpinning of impressive predictive power of AST/ALT ratio with AUROC of 0.83 for advanced fibrosis (F3–F4)(73), leading to AST alone (but not ALT) or AST/ALT ratio being included in multiple non-invasive surrogate liver fibrosis indices such as APRI, FIB-4, NFS, BARD and FibroIndex.(74)

Thus, our findings support OPA1 as a central regulator of the injury-to-scar transition and fibrotic disease severity, independently of steatosis or lipotoxicity. Given that fibrosis stage is the strongest histologic predictor of long-term liver-related outcomes in MASLD,(75) future prospective longitudinal studies are warranted to determine whether OPA1-related variation determines individual fibrosis susceptibility and more rapid progression to advanced fibrosis and end-stage MASH cirrhosis.

Several limitations should be acknowledged. This study provides convergent genetic and tissue expression evidence strongly implicating OPA1 in MASLD fibrosis, but it does not establish causality in human species. Although we demonstrate mtDNA escape into circulation and exaggerated fibrogenic responses in MASH model in hepatocyte-OPA1 deficient mice, species differences in OPA1 biology exist and functional studies in human hepatocytes, Kupffer cells/macrophages, and hepatic stellate cells are needed to unequivocally confirm whether OPA1 dysregulation directly promotes mitochondrial damage signaling, mtDNA release, inflammatory activation, and fibrogenesis in humans. The genetic analyses are also constrained by available GWAS phenotypes, which vary in diagnostic definition, fibrosis staging, ancestry composition, and clinical ascertainment; although MTAG increases discovery power by leveraging shared genetic architecture across related traits,(40) phenotype heterogeneity limits precise inference about the fibrosis stage at which OPA1 is most relevant. In addition, gene-based and regulatory analyses have limited resolution: MAGMA aggregates nearby variant-level signals,(43) and eQTL evidence supports regulatory plausibility but does not prove that OPA1 is the causal effector gene or that regulation occurs in the relevant liver cell type during MASLD progression.(23) Rare-variant and pathway analyses may be underpowered, particularly for stage-specific or ancestry-stratified outcomes, and depend on variant annotation, burden masks, and curated mitochondrial quality-control gene sets. Controls were population-based non-cases rather than participants confirmed free of hepatic steatosis. This may have introduced misclassification from undiagnosed MASLD or liver fibrosis / cirrhosis, likely attenuating associations toward the null and making significant findings more conservative. Finally, transcriptome datasets provides supportive biological context but does not measure OPA1 protein abundance, proteolytic processing, mitochondrial fusion activity, or cristae remodeling, and changes in OPA1 expression in fibrotic MASLD liver may be confounded by changes in cell composition, injury state, sampling depth, or batch effects. Future studies integrating fine-mapping, colocalization, spatial and proteomic profiling, and mechanistic perturbation models will be required to define precisely how OPA1-related mitochondrial quality control contributes to fibrosis progression.

In conclusion, our integrative large-scale human genetic, histopathology analysis and loss-of-function *in vivo* experiments identify OPA1, a central regulator of mitochondrial fusion, as a novel genetic and biological determinant of liver fibrosis in MASLD. Our findings directly link dysregulation of OPA1-dependent mitochondrial turnover to release of pro-fibrogenic mitochondrial DAMPs, triggering hepatic stellate cell activation and liver scarring. These results provide a rationale for mechanistic studies of OPA1-centered mitochondrial quality-control networks and support further investigation of mitochondria-targeted therapeutic strategies for progressive MASLD.

## Acknowledgements

This work was supported by USF Health Graduate Medical Education Research Grant Program awards 2025-26 and 2026-27 to X.Y., National Institutes of Health NIDDK R01DK139288 to Y.V.P., NIH NIGMS grant 5R35GM144103 to H.S. A.H. is supported by the National Institutes of Health (K08DK149008-01), the Sperling Family Foundation Fellowship in Precision Healthcare, the Irving W. and Charlotte F. Rabb Award, and the Beth Israel Deaconess Medical Center Technology Ventures Office Ignition Award. We are grateful to our BIDMC research student interns Nina Ge, Alexander Lugovskoy and Victor Turkowski for their assistance with experiments, sample preparation and data analysis.

