## Supplement for "OPA1 controls mitochondrial dysfunction-driven liver fibrosis in MASLD"

### Correspondence to:

**Table S1. Genome-wide association studies included in the Multi-Trait Analysis of GWAS**

| | Author (Year) | Traits | Ancestry | Sample size | Mean $\chi^2$ | $\lambda$GC | LDSC Intercept (SE) | Attenuation Ratio | Ref. |
|---|---|---|---|---|---|---|---|---|---|
| 1 | Sun Z (2023) | MASLD | European | 32,941 | 1.0574 | 1.0466 | 0.9976 | -0.042 | [1] |
| 2 | Ghodsian N (2021) | MASLD | European | 778,614 | 1.0311 | 1.0285 | 1.0080 | 0.257 | [2] |
| 3 | Fairfield CJ (2021) | MASLD | European | 413,256 | 1.0781 | 1.0679 | 1.0089 | 0.114 | [3] |
| 4 | Kurki MI (2023) | MASLD | European (Finnish) | 500,348 | 1.0749 | 1.0527 | 1.0194 | 0.259 | [4] |
| 5 | Ghouse J (2024) | Cirrhosis of liver | Mixed ethnicity | 3,179,403 | 1.1449 | 1.1082 | 1.0226 | 0.156 | [5] |
| 6 | Ghouse J (2024) | Cirrhosis of liver | European | 2,039,708 | 1.1486 | 1.1144 | 1.0268 | 0.180 | [5] |

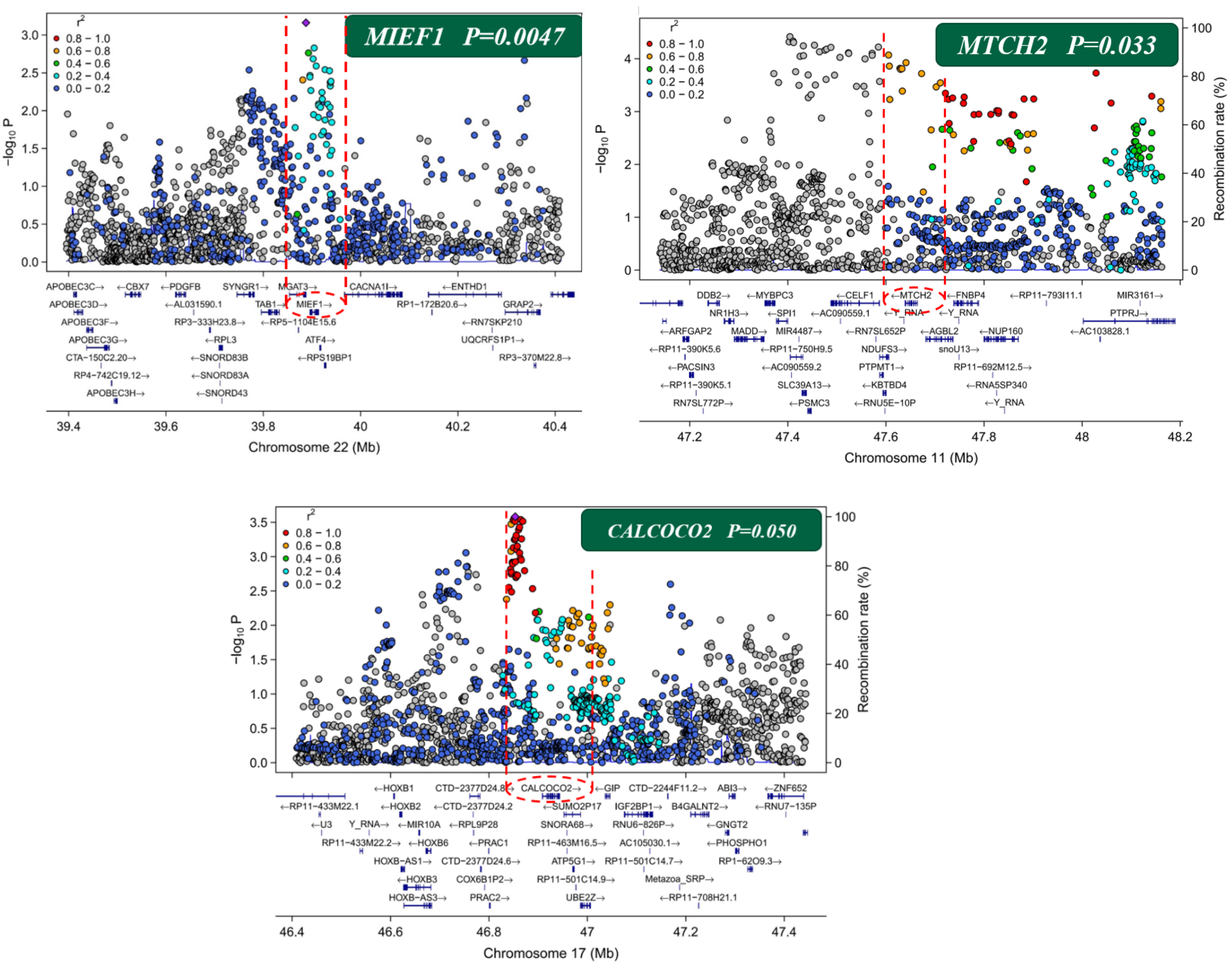


**Figure S1.** Additional mitochondrial genes which demonstrated nominal associations in gene-based association analysis of common variants in MTAG

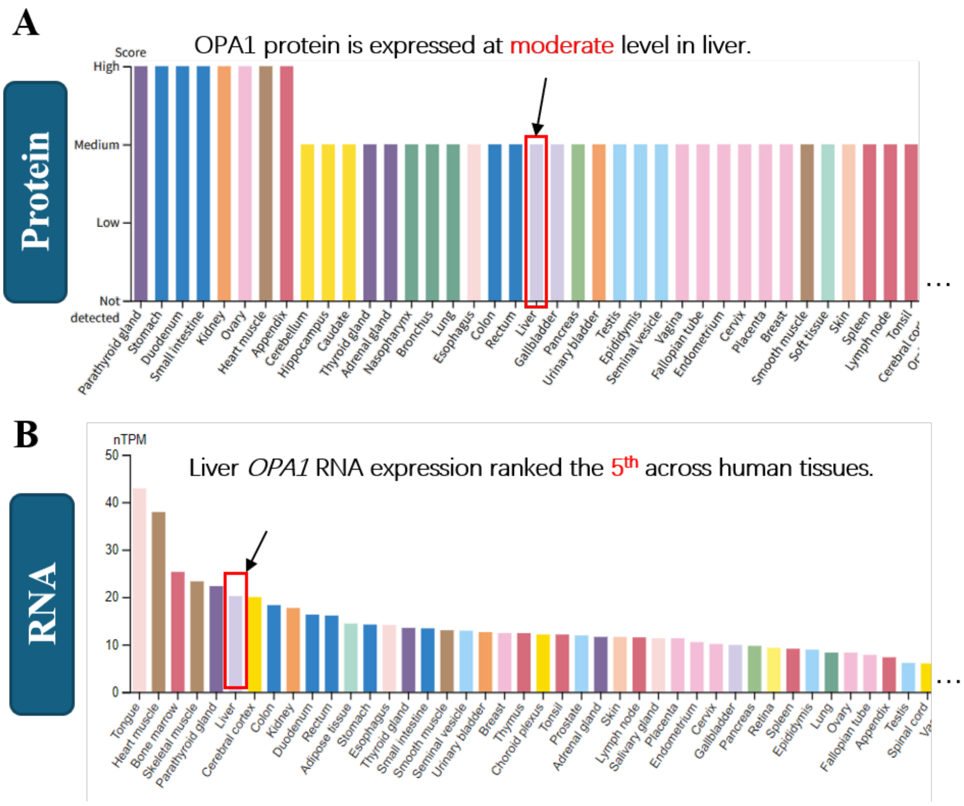


**Figure S2. OPA1 protein and RNA expression across human tissues.** A. The upper panel shows semiquantitative OPA1 protein expression across human tissues, categorized as high, medium, low, or not detected. Liver tissue is highlighted by a red box and arrow, indicating moderate OPA1 protein expression. B. The lower panel shows tissue-level ***OPA1*** RNA expression measured as normalized transcripts per million (nTPM). Liver is highlighted and ranks fifth among the surveyed human tissues for ***OPA1*** RNA expression, supporting prominent hepatic expression of *OPA1* at both the transcript and protein levels.

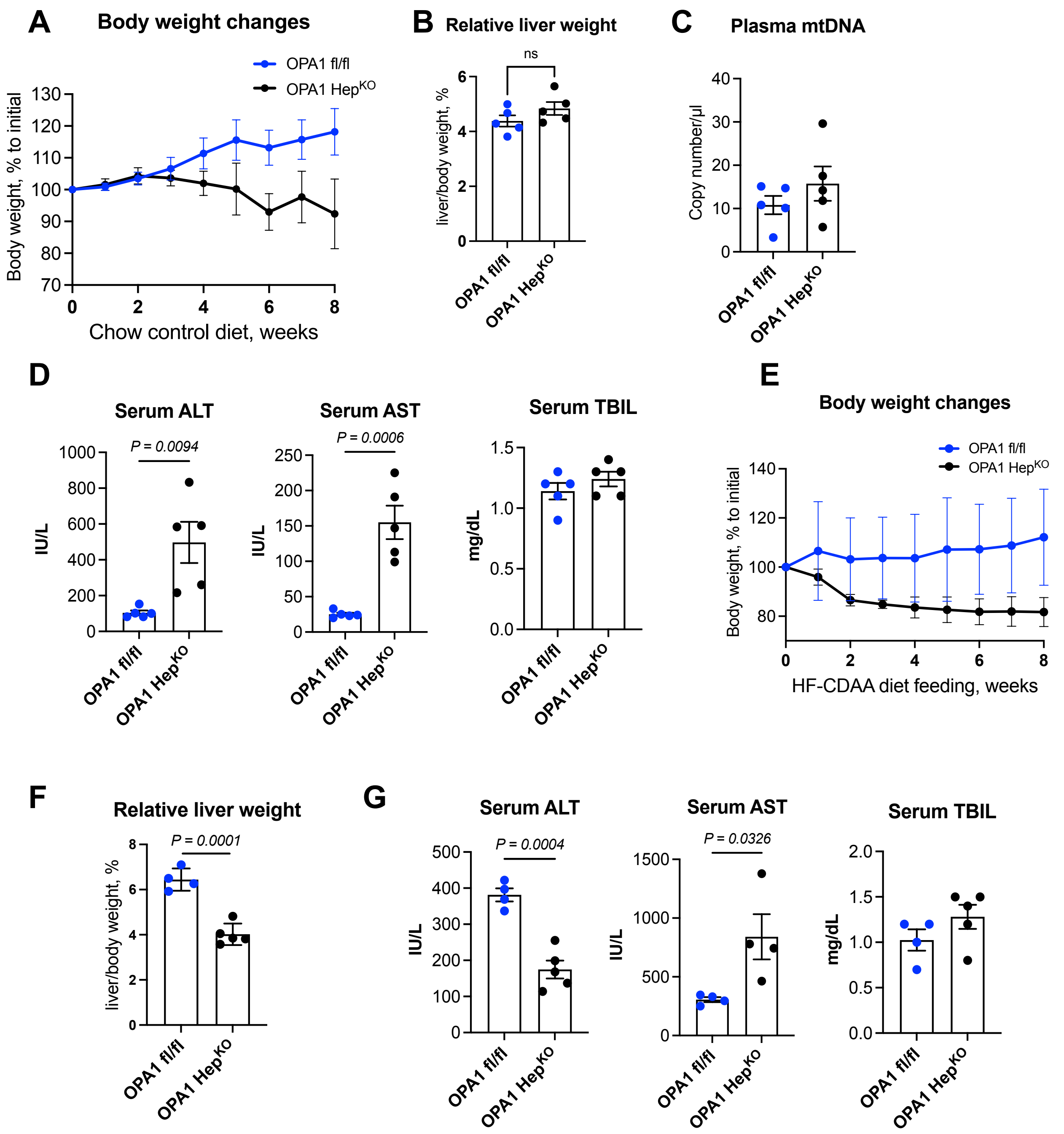


**Figure S3. Body weight changes, hepatomegaly and serum chemistry upon hepatocyte-specific deletion of OPA1 in mice fed control diet and with experimental MASH**. Hepatocyte-specific deletion of OPA1(OPA1 $Hep^{KO}$) was achieved by AAV8-TBG-Cre injection into male floxed OPA1 mice. Equivalent dose of AAV8-EGFP was used in OPA1-sufficient control group ($OPA1^{fl/fl}$). Mice were fed regular chow (A-D, n=5) or MASH diet (HF-CDAA, E-G, n=4-5) for 8 weeks before analysis. (A) Body weight changes, (B) relative liver weight, (C) circulating mtDNA measured via 12S fragment ddPCR and (D) serum ALT, AST and total bilirubin in mice fed control chow diet. (E) Body weight changes, (F) relative liver weight, and (G) serum ALT, AST and total bilirubin in mice fed control chow diet.

**Table S2. Gene burden analysis of loss-of-function and missense variants in liver fibrosis and cirrhosis**

| | | | SKAT-O | SKAT | Burden |
|---|---|---|---|---|---|
| *CRLS1* | OPA1-pathway | LoF | 0. 012 | 0. 008 | 0. 033 |
| *MICU1* | OPA1-interactors | LoF | 0. 012 | 0. 006 | 0. 298 |
| *BAX* | OPA1-pathway | Missense | 0. 013 | 0. 007 | 0. 335 |
| *PMPCA* | OPA1-interactors | Missense | 0. 014 | 1. 000 | 0. 006 |
| *MTX1* | OPA1-pathway | Missense | 0. 025 | 0. 271 | 0. 014 |
| *IMMT* | OPA1-interactors | LoF | 0. 027 | 0. 016 | 0. 240 |
| *IMMT* | OPA1-interactors | LoF | 0. 027 | 0. 016 | 0. 240 |
| *PRELID1* | OPA1-interactors | LoF | 0. 029 | 0. 019 | 0. 097 |
| *MICU1* | OPA1-interactors | Missense | 0. 084 | 0. 336 | 0. 047 |
| *HIGD1A* | OPA1-interactors | LoF | 0. 087 | 0. 063 | 0. 110 |
| *SAMM50* | OPA1-interactors | LoF | 0. 092 | 0. 052 | 0. 386 |
| *YME1L1* | OPA1-interactors | LoF | 0. 115 | 0. 089 | 0. 281 |
| *PMPCB* | OPA1-interactors | Missense | 0. 155 | 0. 855 | 0. 091 |
| *CASP3* | OPA1-pathway | Missense | 0. 165 | 0. 096 | 0. 197 |
| *MICOS13* | OPA1-pathway | Missense | 0. 167 | 0. 097 | 0. 473 |
| *SIRT3* | OPA1-interactors | LoF | 0. 191 | 0. 166 | 0. 130 |
| *CHCHD6* | OPA1-pathway | Missense | 0. 215 | 0. 299 | 0. 131 |
| *APOO* | OPA1-pathway | Missense | 0. 292 | 0. 412 | 0. 185 |
| *YME1L1* | OPA1-interactors | Missense | 0. 314 | 1. 000 | 0. 178 |
| *TAMM41* | OPA1-pathway | LoF | 0. 347 | 0. 211 | 0. 717 |
| *RCC1L* | OPA1-interactors | Missense | 0. 355 | 0. 204 | 0. 485 |
| *TRIAP1* | OPA1-interactors | Missense | 0. 359 | 0. 450 | 0. 243 |
| *HIGD1A* | OPA1-interactors | Missense | 0. 363 | 0. 217 | 0. 472 |
| *AGK* | OPA1-pathway | Missense | 0. 369 | 0. 219 | 0. 429 |
| *PGS1* | OPA1-pathway | Missense | 0. 393 | 0. 281 | 0. 248 |
| *SIRT4* | OPA1-interactors | Missense | 0. 403 | 0. 273 | 0. 513 |
| *AGK* | OPA1-pathway | LoF | 0. 409 | 0. 999 | 0. 251 |
| *PMPCB* | OPA1-interactors | LoF | 0. 418 | 0. 247 | 0. 715 |
| *BAK1* | OPA1-pathway | Missense | 0. 425 | 0. 999 | 0. 257 |
| *APAF1* | OPA1-pathway | Missense | 0. 469 | 0. 296 | 0. 753 |
| *CASP9* | OPA1-pathway | LoF | 0. 501 | 1. 000 | 0. 320 |
| *SAMM50* | OPA1-interactors | Missense | 0. 529 | 0. 593 | 0. 349 |
| *MTX1* | OPA1-pathway | LoF | 0. 575 | 0. 950 | 0. 396 |
| *ATAD3A* | OPA1-interactors | LoF | 0. 585 | 1. 000 | 0. 388 |
| *PGS1* | OPA1-pathway | LoF | 0. 601 | 1. 000 | 0. 393 |
| *PRELID1* | OPA1-interactors | Missense | 0. 645 | 0. 452 | 0. 702 |
| *TOMM70* | OPA1-interactors | Missense | 0. 670 | 0. 535 | 0. 456 |
| *PFKFB3* | OPA1-interactors | Missense | 0. 684 | 0. 511 | 0. 472 |
| *SIRT3* | OPA1-interactors | Missense | 0. 689 | 0. 924 | 0. 496 |
| *CHCHD3* | OPA1-interactors | Missense | 0. 707 | 0. 933 | 0. 489 |
| *CHCHD3* | OPA1-interactors | Missense | 0. 707 | 0. 933 | 0. 489 |
| *CASP3* | OPA1-pathway | LoF | 0. 714 | 0. 906 | 0. 564 |

| | | | SKAT-O | SKAT | Burden |
|---|---|---|---|---|---|
| *CASP9* | OPA1-pathway | Missense | 0.730 | 0.522 | 0.711 |
| *CHCHD6* | OPA1-pathway | LoF | 0.735 | 1.000 | 0.522 |
| *BAK1* | OPA1-pathway | LoF | 0.738 | 1.000 | 0.543 |
| *SIRT4* | OPA1-interactors | LoF | 0.742 | 1.000 | 0.564 |
| *CRLS1* | OPA1-pathway | Missense | 0.748 | 0.555 | 0.586 |
| *PMPCA* | OPA1-interactors | LoF | 0.751 | 1.000 | 0.533 |
| *IMMT* | OPA1-interactors | Missense | 0.759 | 0.577 | 0.875 |
| *IMMT* | OPA1-interactors | Missense | 0.759 | 0.577 | 0.875 |
| *APAF1* | OPA1-pathway | LoF | 0.760 | 0.535 | 0.997 |
| *MTX2* | OPA1-pathway | LoF | 0.809 | 1.000 | 0.613 |
| *MTX2* | OPA1-pathway | Missense | 0.817 | 0.999 | 0.601 |
| *RCC1L* | OPA1-interactors | LoF | 0.818 | 0.603 | 0.940 |
| *MICOS13* | OPA1-pathway | LoF | 0.822 | 1.000 | 0.660 |
| *PFKFB3* | OPA1-interactors | LoF | 0.876 | 1.000 | 0.670 |
| *NEDD4L* | OPA1-interactors | LoF | 0.881 | 1.000 | 0.677 |
| *BAX* | OPA1-pathway | LoF | 0.883 | 1.000 | 0.701 |
| *APOOL* | OPA1-pathway | Missense | 0.886 | 0.852 | 0.688 |
| *TAFAZZIN* | OPA1-pathway | Missense | 0.899 | 0.989 | 0.708 |
| *CHCHD3* | OPA1-interactors | LoF | 0.903 | 1.000 | 0.731 |
| *CHCHD3* | OPA1-interactors | LoF | 0.903 | 1.000 | 0.731 |
| *TOMM70* | OPA1-interactors | LoF | 1.000 | 1.000 | 0.802 |
| *TRIAP1* | OPA1-interactors | LoF | 1.000 | 1.000 | 0.794 |
| *APOO* | OPA1-pathway | LoF | 1.000 | 1.000 | 0.875 |
| *APOOL* | OPA1-pathway | LoF | 1.000 | 1.000 | 0.794 |
| *CYCS* | OPA1-pathway | LoF | 1.000 | 1.000 | 0.888 |
| *MICOS10* | OPA1-pathway | LoF | 1.000 | 0.999 | 0.909 |
| *TAFAZZIN* | OPA1-pathway | LoF | 1.000 | 1.000 | 0.915 |
| *ATAD3A* | OPA1-interactors | Missense | 1.000 | 0.944 | 0.772 |
| *NEDD4L* | OPA1-interactors | Missense | 1.000 | 0.844 | 0.880 |
| *CYCS* | OPA1-pathway | Missense | 1.000 | 0.999 | 0.837 |
| *MICOS10* | OPA1-pathway | Missense | 1.000 | 0.953 | 0.816 |
| *TAMM41* | OPA1-pathway | Missense | 1.000 | 0.953 | 0.928 |

**Table S3. Gene burden analysis of loss-of-function and missense variants in MASLD in UK Biobank**

| | | | SKAT-O | SKAT | Burden |
|---|---|---|---|---|---|
| *BAK1* | OPA1-pathway | LoF | 0. 013 | 0. 013 | 0. 013 |
| *CHCHD6* | OPA1-pathway | LoF | 0. 014 | 0. 008 | 0. 050 |
| *APOOL* | OPA1-pathway | LoF | 0. 017 | 0. 011 | 0. 071 |
| *CYCS* | OPA1-pathway | LoF | 0. 021 | 0. 017 | 0. 036 |
| *CYCS* | OPA1-pathway | Missense | 0. 029 | 0. 019 | 0. 042 |
| *TAFAZZIN* | OPA1-pathway | Missense | 0. 045 | 0. 082 | 0. 030 |
| *TRIAP1* | OPA1-interactors | LoF | 0. 065 | 0. 039 | 0. 207 |
| *PFKFB3* | OPA1-interactors | Missense | 0. 066 | 0. 180 | 0. 037 |
| *IMMT* | OPA1-interactors | Missense | 0. 103 | 0. 197 | 0. 062 |
| *IMMT* | OPA1-interactors | Missense | 0. 103 | 0. 197 | 0. 062 |
| *PMPCA* | OPA1-interactors | Missense | 0. 129 | 0. 148 | 0. 083 |
| *APAF1* | OPA1-pathway | LoF | 0. 157 | 0. 117 | 0. 118 |
| *MICU1* | OPA1-interactors | LoF | 0. 158 | 0. 874 | 0. 092 |
| *PRELID1* | OPA1-interactors | LoF | 0. 162 | 0. 093 | 0. 494 |
| *PFKFB3* | OPA1-interactors | LoF | 0. 181 | 0. 102 | 0. 584 |
| *APOO* | OPA1-pathway | Missense | 0. 186 | 0. 114 | 0. 331 |
| *PMPCB* | OPA1-interactors | LoF | 0. 199 | 0. 109 | 0. 235 |
| *TAMM41* | OPA1-pathway | Missense | 0. 213 | 0. 179 | 0. 127 |
| *BAX* | OPA1-pathway | LoF | 0. 219 | 0. 130 | 0. 427 |
| *AGK* | OPA1-pathway | LoF | 0. 223 | 0. 538 | 0. 130 |
| *HIGD1A* | OPA1-interactors | Missense | 0. 268 | 0. 999 | 0. 154 |
| *HIGD1A* | OPA1-interactors | LoF | 0. 283 | 0. 185 | 0. 720 |
| *MTX1* | OPA1-pathway | Missense | 0. 291 | 0. 217 | 0. 177 |
| *MICOS13* | OPA1-pathway | LoF | 0. 301 | 0. 187 | 0. 667 |
| *APOOL* | OPA1-pathway | Missense | 0. 333 | 0. 194 | 0. 951 |
| *PGS1* | OPA1-pathway | Missense | 0. 335 | 0. 307 | 0. 208 |
| *NEDD4L* | OPA1-interactors | Missense | 0. 340 | 0. 207 | 0. 682 |
| *CASP9* | OPA1-pathway | Missense | 0. 358 | 0. 635 | 0. 219 |
| *CRLS1* | OPA1-pathway | LoF | 0. 397 | 0. 242 | 0. 303 |
| *TAMM41* | OPA1-pathway | LoF | 0. 401 | 0. 244 | 0. 559 |
| *MTX2* | OPA1-pathway | Missense | 0. 417 | 0. 251 | 0. 908 |
| *TRIAP1* | OPA1-interactors | Missense | 0. 420 | 0. 894 | 0. 290 |
| *SAMM50* | OPA1-interactors | Missense | 0. 429 | 0. 399 | 0. 273 |
| *TOMM70* | OPA1-interactors | Missense | 0. 435 | 0. 267 | 0. 786 |
| *YME1L1* | OPA1-interactors | LoF | 0. 481 | 0. 376 | 0. 373 |
| *CRLS1* | OPA1-pathway | Missense | 0. 497 | 0. 372 | 0. 329 |
| *AGK* | OPA1-pathway | Missense | 0. 507 | 0. 314 | 0. 550 |
| *SIRT3* | OPA1-interactors | LoF | 0. 522 | 0. 832 | 0. 382 |
| *MTX2* | OPA1-pathway | LoF | 0. 558 | 1. 000 | 0. 369 |
| *APAF1* | OPA1-pathway | Missense | 0. 598 | 0. 397 | 0. 529 |
| *IMMT* | OPA1-interactors | LoF | 0. 626 | 0. 401 | 0. 841 |
| *IMMT* | OPA1-interactors | LoF | 0. 626 | 0. 401 | 0. 841 |
| *MICOS10* | OPA1-pathway | Missense | 0. 664 | 0. 482 | 0. 720 |
| *MICU1* | OPA1-interactors | Missense | 0. 689 | 0. 471 | 0. 652 |

| | | | SKAT-O | SKAT | Burden |
|---|---|---|---|---|---|
| *NEDD4L* | OPA1-interactors | LoF | 0.697 | 1.000 | 0.468 |
| *CHCHD3* | OPA1-interactors | LoF | 0.749 | 1.000 | 0.539 |
| *CHCHD3* | OPA1-interactors | LoF | 0.749 | 1.000 | 0.539 |
| *SIRT3* | OPA1-interactors | Missense | 0.751 | 0.691 | 0.558 |
| *CASP9* | OPA1-pathway | LoF | 0.792 | 0.848 | 0.589 |
| *RCC1L* | OPA1-interactors | LoF | 0.829 | 0.978 | 0.616 |
| *TOMM70* | OPA1-interactors | LoF | 0.842 | 1.000 | 0.641 |
| *MTX1* | OPA1-pathway | LoF | 0.860 | 0.715 | 0.897 |
| *SAMM50* | OPA1-interactors | LoF | 0.862 | 0.661 | 0.866 |
| *CHCHD3* | OPA1-interactors | Missense | 0.869 | 0.670 | 0.982 |
| *CHCHD3* | OPA1-interactors | Missense | 0.869 | 0.670 | 0.982 |
| *CASP3* | OPA1-pathway | Missense | 0.872 | 0.803 | 0.695 |
| *SIRT4* | OPA1-interactors | LoF | 0.879 | 0.725 | 0.955 |
| *BAX* | OPA1-pathway | Missense | 0.885 | 0.722 | 0.699 |
| *PMPCA* | OPA1-interactors | LoF | 0.898 | 0.894 | 0.727 |
| *PMPCB* | OPA1-interactors | Missense | 0.902 | 0.747 | 0.775 |
| *BAK1* | OPA1-pathway | Missense | 0.907 | 0.773 | 0.729 |
| *APOO* | OPA1-pathway | LoF | 0.913 | 1.000 | 0.765 |
| *RCC1L* | OPA1-interactors | Missense | 0.932 | 0.756 | 0.790 |
| *YME1L1* | OPA1-interactors | Missense | 0.936 | 0.767 | 0.959 |
| *ATAD3A* | OPA1-interactors | LoF | 1.000 | 0.898 | 0.820 |
| *CASP3* | OPA1-pathway | LoF | 1.000 | 0.922 | 0.834 |
| *MICOS10* | OPA1-pathway | LoF | 1.000 | 1.000 | 0.835 |
| *PGS1* | OPA1-pathway | LoF | 1.000 | 0.902 | 0.942 |
| *TAFAZZIN* | OPA1-pathway | LoF | 1.000 | 1.000 | 0.861 |
| *ATAD3A* | OPA1-interactors | Missense | 1.000 | 0.938 | 0.819 |
| *PRELID1* | OPA1-interactors | Missense | 1.000 | 0.989 | 0.819 |
| *SIRT4* | OPA1-interactors | Missense | 1.000 | 0.991 | 0.832 |
| *CHCHD6* | OPA1-pathway | Missense | 1.000 | 0.934 | 0.995 |
| *MICOS13* | OPA1-pathway | Missense | 1.000 | 0.807 | 0.881 |

**Table S4. *OPA1* eQTL summary across different tissues**

| MTAG SNP | eQTL SNP | Position GRCH37 | $R^2$ | Tissue | Effect AF | Effect_Size | $P_{eQTL}$ | a1 | a2 | $N_{MTAG}$ | a1 frequency | Beta | $P_{MTAG}$ |
|---|---|---|---|---|---|---|---|---|---|---|---|---|---|
| rs78912308 | rs78912308 | chr3:193266100 | 1.00 | Cells - Cultured fibroblasts | A=0.101 | -0.2473 | 6.77E-23 | C | A | 1621580 | 0.90 | -0.00409 | 9.87E-03 |
| rs78912308 | rs78912308 | chr3:193266100 | 1.00 | Whole Blood | A=0.101 | -0.1833 | 9.55E-07 | C | A | 1621580 | 0.90 | -0.00409 | 9.87E-03 |
| rs76205181 | rs78912308 | chr3:193273067 | 0.94 | Whole Blood | T=0.096 | -0.1787 | 1.70E-06 | C | T | 1522690 | 0.90 | -0.0053 | 1.11E-03 |
| rs76205181 | rs78912308 | chr3:193273067 | 0.94 | Cells - Cultured fibroblasts | T=0.096 | -0.2439 | 2.19E-22 | C | T | 1522690 | 0.90 | -0.0053 | 1.11E-03 |
| rs1872532 | rs1872532 | chr3:193284624 | 1.00 | Whole Blood | C=0.328 | -0.0913 | 6.62E-06 | T | C | 1800310 | 0.67 | -0.00276 | 3.94E-03 |
| rs1872532 | rs1872532 | chr3:193284624 | 1.00 | Cells - Cultured fibroblasts | C=0.328 | -0.0739 | 1.09E-06 | T | C | 1800310 | 0.67 | -0.00276 | 3.94E-03 |
| rs7625424 | rs1872532 | chr3:193285897 | 1.00 | Cells - Cultured fibroblasts | C=0.328 | -0.0744 | 1.06E-06 | T | C | 1800310 | 0.66 | -0.0027 | 4.67E-03 |
| rs7625424 | rs1872532 | chr3:193285897 | 1.00 | Whole Blood | C=0.328 | -0.0872 | 1.68E-05 | T | C | 1800310 | 0.66 | -0.0027 | 4.67E-03 |
| rs6444728 | rs1872532 | chr3:193286182 | 1.00 | Cells - Cultured fibroblasts | T=0.328 | -0.0744 | 1.06E-06 | C | T | 1624360 | 0.66 | -0.00332 | 8.54E-04 |
| rs6444728 | rs1872532 | chr3:193286182 | 1.00 | Whole Blood | T=0.328 | -0.0872 | 1.68E-05 | C | T | 1624360 | 0.66 | -0.00332 | 8.54E-04 |
| rs9869967 | rs1872532 | chr3:193287410 | 0.87 | Whole Blood | A=0.298 | -0.0896 | 1.79E-05 | G | A | 1701420 | 0.70 | -0.00327 | 1.19E-03 |
| rs9869967 | rs1872532 | chr3:193287410 | 0.87 | Cells - Cultured fibroblasts | A=0.298 | -0.0923 | 1.97E-09 | G | A | 1701420 | 0.70 | -0.00327 | 1.19E-03 |
| rs13326186 | rs1872532 | chr3:193288057 | 0.87 | Cells - Cultured fibroblasts | A=0.298 | -0.0928 | 1.55E-09 | G | A | 1800310 | 0.70 | -0.00289 | 3.26E-03 |
| rs13326186 | rs1872532 | chr3:193288057 | 0.87 | Whole Blood | A=0.298 | -0.0897 | 1.72E-05 | G | A | 1800310 | 0.70 | -0.00289 | 3.26E-03 |
| rs13326189 | rs1872532 | chr3:193288133 | 1.00 | Cells - Cultured fibroblasts | A=0.328 | -0.0744 | 1.06E-06 | G | A | 1800310 | 0.66 | -0.00269 | 4.90E-03 |
| rs13326189 | rs1872532 | chr3:193288133 | 1.00 | Whole Blood | A=0.328 | -0.0872 | 1.68E-05 | G | A | 1800310 | 0.66 | -0.00269 | 4.90E-03 |
| rs3863983 | rs1872532 | chr3:193288804 | 1.00 | Whole Blood | C=0.328 | -0.0864 | 2.16E-05 | T | C | 1624360 | 0.66 | -0.00332 | 8.75E-04 |
| rs3863983 | rs1872532 | chr3:193288804 | 1.00 | Cells - Cultured fibroblasts | C=0.328 | -0.0739 | 1.33E-06 | T | C | 1624360 | 0.66 | -0.00332 | 8.75E-04 |
| rs7650397 | rs1872532 | chr3:193290121 | 0.93 | Cells - Cultured fibroblasts | C=0.313 | -0.0778 | 4.44E-07 | T | C | 1800310 | 0.67 | -0.00276 | 4.09E-03 |
| rs7650397 | rs1872532 | chr3:193290121 | 0.93 | Whole Blood | C=0.313 | -0.0885 | 1.41E-05 | T | C | 1800310 | 0.67 | -0.00276 | 4.09E-03 |
| rs11922377 | rs1872532 | chr3:193290542 | 0.93 | Whole Blood | G=0.313 | -0.0885 | 1.41E-05 | A | G | 1645020 | 0.67 | -0.00286 | 3.84E-03 |
| rs11922377 | rs1872532 | chr3:193290542 | 0.93 | Cells - Cultured fibroblasts | G=0.313 | -0.0778 | 4.44E-07 | A | G | 1645020 | 0.67 | -0.00286 | 3.84E-03 |
| rs4597663 | rs4597663 | chr3:193291405 | 1.00 | Whole Blood | T=0.47 | -0.0988 | 1.51E-07 | C | T | 1800310 | 0.52 | -0.0029 | 1.36E-03 |
| rs4597663 | rs2935303 | chr3:193291405 | 0.70 | Whole Blood | T=0.47 | -0.0988 | 1.51E-07 | C | T | 1800310 | 0.52 | -0.0029 | 1.36E-03 |
| rs13067143 | rs2935303 | chr3:193291640 | 0.70 | Whole Blood | T=0.47 | -0.0988 | 1.51E-07 | G | T | 1701420 | 0.52 | -0.00356 | 1.26E-04 |
| rs13067143 | rs4597663 | chr3:193291640 | 1.00 | Whole Blood | T=0.47 | -0.0988 | 1.51E-07 | G | T | 1701420 | 0.52 | -0.00356 | 1.26E-04 |
| rs11916762 | rs4597663 | chr3:193292035 | 1.00 | Whole Blood | A=0.47 | -0.0988 | 1.51E-07 | G | A | 1800310 | 0.52 | -0.00291 | 1.31E-03 |
| rs11916762 | rs2935303 | chr3:193292035 | 0.70 | Whole Blood | A=0.47 | -0.0988 | 1.51E-07 | G | A | 1800310 | 0.52 | -0.00291 | 1.31E-03 |

| MTAG SNP | eQTL SNP | Position GRCH37 | $R^2$ | Tissue | Effect AF | Effect_Size | $P_{eQTL}$ | a1 | a2 | $N_{MTAG}$ | a1 frequency | Beta | $P_{MTAG}$ |
|---|---|---|---|---|---|---|---|---|---|---|---|---|---|
| rs56102513 | rs78912308 | chr3:193292325 | 0.94 | Cells - Cultured fibroblasts | C=0.096 | -0.2395 | 4.92E-22 | T | C | 1621580 | 0.90 | -0.0043 | 6.73E-03 |
| rs56102513 | rs78912308 | chr3:193292325 | 0.94 | Pancreas | C=0.096 | -0.3023 | 3.87E-05 | T | C | 1621580 | 0.90 | -0.0043 | 6.73E-03 |
| rs56102513 | rs78912308 | chr3:193292325 | 0.94 | Whole Blood | C=0.096 | -0.1840 | 4.85E-07 | T | C | 1621580 | 0.90 | -0.0043 | 6.73E-03 |
| rs7615623 | rs4597663 | chr3:193293157 | 0.88 | Cells - Cultured fibroblasts | C=0.439 | -0.0680 | 3.06E-06 | T | C | 1800310 | 0.56 | -0.00296 | 1.19E-03 |
| rs7615623 | rs4597663 | chr3:193293157 | 0.88 | Whole Blood | C=0.439 | -0.0977 | 2.47E-07 | T | C | 1800310 | 0.56 | -0.00296 | 1.19E-03 |
| rs7626230 | rs4597663 | chr3:193293533 | 1.00 | Whole Blood | C=0.47 | -0.0988 | 1.51E-07 | A | C | 1701420 | 0.52 | -0.00353 | 1.43E-04 |
| rs7626230 | rs2935303 | chr3:193293533 | 0.70 | Whole Blood | C=0.47 | -0.0988 | 1.51E-07 | A | C | 1701420 | 0.52 | -0.00353 | 1.43E-04 |
| rs12696669 | rs4597663 | chr3:193293699 | 1.00 | Whole Blood | A=0.47 | -0.0988 | 1.51E-07 | C | A | 1800310 | 0.52 | -0.0029 | 1.40E-03 |
| rs12696669 | rs2935303 | chr3:193293699 | 0.70 | Whole Blood | A=0.47 | -0.0988 | 1.51E-07 | C | A | 1800310 | 0.52 | -0.0029 | 1.40E-03 |
| rs13324968 | rs4597663 | chr3:193293922 | 0.88 | Cells - Cultured fibroblasts | G=0.439 | -0.0687 | 2.36E-06 | A | G | 1800310 | 0.56 | -0.00294 | 1.28E-03 |
| rs13324968 | rs4597663 | chr3:193293922 | 0.88 | Whole Blood | G=0.439 | -0.0980 | 2.21E-07 | A | G | 1800310 | 0.56 | -0.00294 | 1.28E-03 |
| rs61289905 | rs4597663 | chr3:193295208 | 0.88 | Cells - Cultured fibroblasts | C=0.439 | -0.0680 | 3.06E-06 | T | C | 1800310 | 0.56 | -0.00297 | 1.13E-03 |
| rs61289905 | rs4597663 | chr3:193295208 | 0.88 | Whole Blood | C=0.439 | -0.0977 | 2.47E-07 | T | C | 1800310 | 0.56 | -0.00297 | 1.13E-03 |
| rs10804956 | rs4597663 | chr3:193297321 | 1.00 | Whole Blood | C=0.47 | -0.0990 | 1.46E-07 | T | C | 1800310 | 0.52 | -0.00291 | 1.33E-03 |
| rs10804956 | rs2935303 | chr3:193297321 | 0.70 | Whole Blood | C=0.47 | -0.0990 | 1.46E-07 | T | C | 1800310 | 0.52 | -0.00291 | 1.33E-03 |
| rs77568410 | rs78912308 | chr3:193297467 | 0.94 | Pancreas | T=0.096 | -0.3023 | 3.87E-05 | C | T | 1621580 | 0.90 | -0.00432 | 6.48E-03 |
| rs77568410 | rs78912308 | chr3:193297467 | 0.94 | Whole Blood | T=0.096 | -0.1840 | 4.85E-07 | C | T | 1621580 | 0.90 | -0.00432 | 6.48E-03 |
| rs77568410 | rs78912308 | chr3:193297467 | 0.94 | Cells - Cultured fibroblasts | T=0.096 | -0.2395 | 4.92E-22 | C | T | 1621580 | 0.90 | -0.00432 | 6.48E-03 |
| rs75745995 | rs78912308 | chr3:193298153 | 0.94 | Cells - Cultured fibroblasts | A=0.096 | -0.2395 | 4.92E-22 | G | A | 1466300 | 0.90 | -0.00451 | 5.95E-03 |
| rs75745995 | rs78912308 | chr3:193298153 | 0.94 | Pancreas | A=0.096 | -0.3023 | 3.87E-05 | G | A | 1466300 | 0.90 | -0.00451 | 5.95E-03 |
| rs75745995 | rs78912308 | chr3:193298153 | 0.94 | Whole Blood | A=0.096 | -0.1840 | 4.85E-07 | G | A | 1466300 | 0.90 | -0.00451 | 5.95E-03 |
| rs6444729 | rs2935303 | chr3:193298641 | 0.70 | Whole Blood | C=0.47 | -0.0990 | 1.46E-07 | T | C | 1800310 | 0.52 | -0.0029 | 1.36E-03 |
| rs6444729 | rs4597663 | chr3:193298641 | 1.00 | Whole Blood | C=0.47 | -0.0990 | 1.46E-07 | T | C | 1800310 | 0.52 | -0.0029 | 1.36E-03 |
| rs2134880 | rs2935303 | chr3:193299022 | 0.70 | Whole Blood | C=0.47 | -0.0990 | 1.46E-07 | A | C | 1800310 | 0.52 | -0.0029 | 1.39E-03 |
| rs2134880 | rs4597663 | chr3:193299022 | 1.00 | Whole Blood | C=0.47 | -0.0990 | 1.46E-07 | A | C | 1800310 | 0.52 | -0.0029 | 1.39E-03 |
| rs2134881 | rs4597663 | chr3:193299030 | 1.00 | Whole Blood | C=0.47 | -0.0990 | 1.46E-07 | T | C | 1800310 | 0.52 | -0.00289 | 1.44E-03 |
| rs2134881 | rs2935303 | chr3:193299030 | 0.70 | Whole Blood | C=0.47 | -0.0990 | 1.46E-07 | T | C | 1800310 | 0.52 | -0.00289 | 1.44E-03 |
| rs2174320 | rs2935303 | chr3:193299124 | 0.70 | Whole Blood | A=0.47 | -0.0990 | 1.46E-07 | G | A | 1800310 | 0.52 | -0.0029 | 1.37E-03 |
| rs2174320 | rs4597663 | chr3:193299124 | 1.00 | Whole Blood | A=0.47 | -0.0990 | 1.46E-07 | G | A | 1800310 | 0.52 | -0.0029 | 1.37E-03 |
| rs2174321 | rs4597663 | chr3:193299155 | 0.88 | Cells - Cultured fibroblasts | A=0.439 | -0.0680 | 3.06E-06 | G | A | 1701420 | 0.56 | -0.00351 | 1.71E-04 |

| MTAG SNP | eQTL SNP | Position GRCH37 | $R^2$ | Tissue | Effect AF | Effect_Size | $P_{eQTL}$ | a1 | a2 | $N_{MTAG}$ | a1 frequency | Beta | $P_{MTAG}$ |
|---|---|---|---|---|---|---|---|---|---|---|---|---|---|
| rs2174321 | rs4597663 | chr3:193299155 | 0.88 | Whole Blood | A=0.439 | -0.0977 | 2.47E-07 | G | A | 1701420 | 0.56 | -0.00351 | 1.71E-04 |
| rs7650353 | rs2935303 | chr3:193299182 | 0.70 | Whole Blood | T=0.47 | -0.0990 | 1.46E-07 | C | T | 1800310 | 0.52 | -0.00289 | 1.43E-03 |
| rs7650353 | rs4597663 | chr3:193299182 | 1.00 | Whole Blood | T=0.47 | -0.0990 | 1.46E-07 | C | T | 1800310 | 0.52 | -0.00289 | 1.43E-03 |
| rs56002867 | rs78912308 | chr3:193299226 | 0.94 | Cells - Cultured fibroblasts | T=0.096 | -0.2395 | 4.92E-22 | C | T | 1621580 | 0.90 | -0.00426 | 7.23E-03 |
| rs56002867 | rs78912308 | chr3:193299226 | 0.94 | Whole Blood | T=0.096 | -0.1840 | 4.85E-07 | C | T | 1621580 | 0.90 | -0.00426 | 7.23E-03 |
| rs56002867 | rs78912308 | chr3:193299226 | 0.94 | Pancreas | T=0.096 | -0.3023 | 3.87E-05 | C | T | 1621580 | 0.90 | -0.00426 | 7.23E-03 |
| rs7650444 | rs4597663 | chr3:193299273 | 1.00 | Whole Blood | T=0.47 | -0.0990 | 1.46E-07 | C | T | 1645020 | 0.52 | -0.00294 | 1.68E-03 |
| rs7650444 | rs2935303 | chr3:193299273 | 0.70 | Whole Blood | T=0.47 | -0.0990 | 1.46E-07 | C | T | 1645020 | 0.52 | -0.00294 | 1.68E-03 |
| rs7628243 | rs4597663 | chr3:193299325 | 1.00 | Whole Blood | G=0.47 | -0.0990 | 1.46E-07 | A | G | 1800310 | 0.52 | -0.00289 | 1.45E-03 |
| rs7628243 | rs2935303 | chr3:193299325 | 0.70 | Whole Blood | G=0.47 | -0.0990 | 1.46E-07 | A | G | 1800310 | 0.52 | -0.00289 | 1.45E-03 |
| rs75547111 | rs78912308 | chr3:193299411 | 0.94 | Cells - Cultured fibroblasts | C=0.096 | -0.2395 | 4.92E-22 | T | C | 1800310 | 0.90 | -0.00391 | 9.24E-03 |
| rs75547111 | rs78912308 | chr3:193299411 | 0.94 | Pancreas | C=0.096 | -0.3023 | 3.87E-05 | T | C | 1800310 | 0.90 | -0.00391 | 9.24E-03 |
| rs75547111 | rs78912308 | chr3:193299411 | 0.94 | Whole Blood | C=0.096 | -0.1840 | 4.85E-07 | T | C | 1800310 | 0.90 | -0.00391 | 9.24E-03 |
| rs7620342 | rs2935303 | chr3:193299614 | 0.70 | Whole Blood | G=0.47 | -0.0990 | 1.46E-07 | T | G | 1800310 | 0.52 | -0.00291 | 1.33E-03 |
| rs7620342 | rs4597663 | chr3:193299614 | 1.00 | Whole Blood | G=0.47 | -0.0990 | 1.46E-07 | T | G | 1800310 | 0.52 | -0.00291 | 1.33E-03 |
| rs7653046 | rs4597663 | chr3:193299631 | 0.88 | Whole Blood | T=0.439 | -0.0977 | 2.47E-07 | C | T | 1800310 | 0.56 | -0.00293 | 1.31E-03 |
| rs7653046 | rs4597663 | chr3:193299631 | 0.88 | Cells - Cultured fibroblasts | T=0.439 | -0.0680 | 3.06E-06 | C | T | 1800310 | 0.56 | -0.00293 | 1.31E-03 |
| rs10937592 | rs2935303 | chr3:193301038 | 0.68 | Whole Blood | A=0.475 | -0.0982 | 1.78E-07 | G | A | 1624360 | 0.52 | -0.00372 | 7.97E-05 |
| rs10937592 | rs4597663 | chr3:193301038 | 0.98 | Whole Blood | A=0.475 | -0.0982 | 1.78E-07 | G | A | 1624360 | 0.52 | -0.00372 | 7.97E-05 |
| rs7634678 | rs2935303 | chr3:193301531 | 0.70 | Whole Blood | G=0.47 | -0.0990 | 1.46E-07 | A | G | 1213560 | 0.49 | -0.00351 | 9.78E-04 |
| rs7634678 | rs4597663 | chr3:193301531 | 1.00 | Whole Blood | G=0.47 | -0.0990 | 1.46E-07 | A | G | 1213560 | 0.49 | -0.00351 | 9.78E-04 |
| rs7627228 | rs2935303 | chr3:193302226 | 0.70 | Whole Blood | G=0.47 | -0.0990 | 1.46E-07 | T | G | 1800310 | 0.52 | -0.00288 | 1.49E-03 |
| rs7627228 | rs4597663 | chr3:193302226 | 1.00 | Whole Blood | G=0.47 | -0.0990 | 1.46E-07 | T | G | 1800310 | 0.52 | -0.00288 | 1.49E-03 |
| rs9846719 | rs2935303 | chr3:193302720 | 0.70 | Whole Blood | T=0.47 | -0.0985 | 1.59E-07 | C | T | 1800310 | 0.53 | -0.00278 | 2.13E-03 |
| rs9846719 | rs4597663 | chr3:193302720 | 1.00 | Whole Blood | T=0.47 | -0.0985 | 1.59E-07 | C | T | 1800310 | 0.53 | -0.00278 | 2.13E-03 |
| rs34023161 | rs2935303 | chr3:193302752 | 0.70 | Whole Blood | T=0.47 | -0.0985 | 1.59E-07 | C | T | 1392290 | 0.53 | -0.00357 | 3.34E-04 |
| rs34023161 | rs4597663 | chr3:193302752 | 1.00 | Whole Blood | T=0.47 | -0.0985 | 1.59E-07 | C | T | 1392290 | 0.53 | -0.00357 | 3.34E-04 |
| rs7618459 | rs4597663 | chr3:193303032 | 0.88 | Whole Blood | T=0.439 | -0.0977 | 2.47E-07 | C | T | 1701420 | 0.56 | -0.00351 | 1.72E-04 |
| rs7618459 | rs4597663 | chr3:193303032 | 0.88 | Cells - Cultured fibroblasts | T=0.439 | -0.0680 | 3.06E-06 | C | T | 1701420 | 0.56 | -0.00351 | 1.72E-04 |
| rs7618541 | rs4597663 | chr3:193303101 | 0.88 | Cells - Cultured fibroblasts | T=0.439 | -0.0683 | 2.92E-06 | C | T | 1723250 | 0.55 | -0.00318 | 5.74E-04 |

| MTAG SNP | eQTL SNP | Position GRCH37 | $R^2$ | Tissue | Effect AF | Effect_Size | $P_{eQTL}$ | a1 | a2 | $N_{MTAG}$ | a1 frequency | Beta | $P_{MTAG}$ |
|---|---|---|---|---|---|---|---|---|---|---|---|---|---|
| rs7618541 | rs4597663 | chr3:193303101 | 0.88 | Whole Blood | T=0.439 | -0.0980 | 2.34E-07 | C | T | 1723250 | 0.55 | -0.00318 | 5.74E-04 |
| rs13064790 | rs4597663 | chr3:193303535 | 0.88 | Whole Blood | A=0.439 | -0.0987 | 1.88E-07 | G | A | 1800310 | 0.56 | -0.00297 | 1.11E-03 |
| rs13064790 | rs4597663 | chr3:193303535 | 0.88 | Cells - Cultured fibroblasts | A=0.439 | -0.0678 | 3.58E-06 | G | A | 1800310 | 0.56 | -0.00297 | 1.11E-03 |
| rs9856156 | rs4597663 | chr3:193314595 | 1.00 | Whole Blood | A=0.47 | -0.1020 | 6.64E-08 | G | A | 1701420 | 0.53 | -0.00363 | 9.26E-05 |
| rs9856156 | rs2935303 | chr3:193314595 | 0.70 | Cells - Cultured fibroblasts | A=0.47 | -0.0586 | 7.01E-05 | G | A | 1701420 | 0.53 | -0.00363 | 9.26E-05 |
| rs9856156 | rs4597663 | chr3:193314595 | 1.00 | Cells - Cultured fibroblasts | A=0.47 | -0.0586 | 7.01E-05 | G | A | 1701420 | 0.53 | -0.00363 | 9.26E-05 |
| rs9856156 | rs2935303 | chr3:193314595 | 0.70 | Whole Blood | A=0.47 | -0.1020 | 6.64E-08 | G | A | 1701420 | 0.53 | -0.00363 | 9.26E-05 |
| rs75977202 | rs78912308 | chr3:193315176 | 0.94 | Cells - Cultured fibroblasts | A=0.096 | -0.2407 | 2.90E-22 | G | A | 1621580 | 0.90 | -0.00455 | 4.23E-03 |
| rs75977202 | rs78912308 | chr3:193315176 | 0.94 | Whole Blood | A=0.096 | -0.1849 | 4.79E-07 | G | A | 1621580 | 0.90 | -0.00455 | 4.23E-03 |
| rs75977202 | rs78912308 | chr3:193315176 | 0.94 | Pancreas | A=0.096 | -0.3050 | 3.79E-05 | G | A | 1621580 | 0.90 | -0.00455 | 4.23E-03 |
| rs4263236 | rs4597663 | chr3:193315331 | 1.00 | Cells - Cultured fibroblasts | T=0.47 | -0.0607 | 4.86E-05 | G | T | 1800310 | 0.53 | -0.00301 | 9.02E-04 |
| rs4263236 | rs2935303 | chr3:193315331 | 0.70 | Whole Blood | T=0.47 | -0.1006 | 1.08E-07 | G | T | 1800310 | 0.53 | -0.00301 | 9.02E-04 |
| rs4263236 | rs4597663 | chr3:193315331 | 1.00 | Whole Blood | T=0.47 | -0.1006 | 1.08E-07 | G | T | 1800310 | 0.53 | -0.00301 | 9.02E-04 |
| rs4263236 | rs2935303 | chr3:193315331 | 0.70 | Cells - Cultured fibroblasts | T=0.47 | -0.0607 | 4.86E-05 | G | T | 1800310 | 0.53 | -0.00301 | 9.02E-04 |
| rs12696670 | rs4597663 | chr3:193316048 | 0.88 | Cells - Cultured fibroblasts | G=0.439 | -0.0676 | 3.62E-06 | A | G | 1800310 | 0.56 | -0.00291 | 1.43E-03 |
| rs12696670 | rs4597663 | chr3:193316048 | 0.88 | Whole Blood | G=0.439 | -0.0973 | 2.51E-07 | A | G | 1800310 | 0.56 | -0.00291 | 1.43E-03 |
| rs6800240 | rs2935303 | chr3:193318250 | 0.70 | Whole Blood | C=0.47 | -0.1036 | 4.26E-08 | T | C | 1800310 | 0.52 | -0.00291 | 1.31E-03 |
| rs6800240 | rs2935303 | chr3:193318250 | 0.70 | Cells - Cultured fibroblasts | C=0.47 | -0.0606 | 4.30E-05 | T | C | 1800310 | 0.52 | -0.00291 | 1.31E-03 |
| rs6800240 | rs4597663 | chr3:193318250 | 1.00 | Whole Blood | C=0.47 | -0.1036 | 4.26E-08 | T | C | 1800310 | 0.52 | -0.00291 | 1.31E-03 |
| rs6800240 | rs4597663 | chr3:193318250 | 1.00 | Cells - Cultured fibroblasts | C=0.47 | -0.0606 | 4.30E-05 | T | C | 1800310 | 0.52 | -0.00291 | 1.31E-03 |
| rs6790120 | rs4597663 | chr3:193318576 | 0.88 | Whole Blood | T=0.439 | -0.0973 | 2.51E-07 | C | T | 1800310 | 0.56 | -0.00292 | 1.34E-03 |
| rs6790120 | rs4597663 | chr3:193318576 | 0.88 | Cells - Cultured fibroblasts | T=0.439 | -0.0676 | 3.62E-06 | C | T | 1800310 | 0.56 | -0.00292 | 1.34E-03 |
| rs6794338 | rs2935303 | chr3:193320149 | 0.70 | Whole Blood | A=0.47 | -0.1036 | 4.26E-08 | G | A | 1701420 | 0.53 | -0.00361 | 1.02E-04 |
| rs6794338 | rs4597663 | chr3:193320149 | 1.00 | Whole Blood | A=0.47 | -0.1036 | 4.26E-08 | G | A | 1701420 | 0.53 | -0.00361 | 1.02E-04 |
| rs6794338 | rs4597663 | chr3:193320149 | 1.00 | Cells - Cultured fibroblasts | A=0.47 | -0.0606 | 4.30E-05 | G | A | 1701420 | 0.53 | -0.00361 | 1.02E-04 |
| rs6794338 | rs2935303 | chr3:193320149 | 0.70 | Cells - Cultured fibroblasts | A=0.47 | -0.0606 | 4.30E-05 | G | A | 1701420 | 0.53 | -0.00361 | 1.02E-04 |
| rs7619129 | rs4597663 | chr3:193320736 | 0.88 | Cells - Cultured fibroblasts | G=0.439 | -0.0676 | 3.62E-06 | A | G | 1701420 | 0.56 | -0.00351 | 1.71E-04 |
| rs7619129 | rs4597663 | chr3:193320736 | 0.88 | Whole Blood | G=0.439 | -0.0973 | 2.51E-07 | A | G | 1701420 | 0.56 | -0.00351 | 1.71E-04 |
| rs11925699 | rs2935303 | chr3:193321115 | 0.70 | Whole Blood | A=0.47 | -0.1059 | 2.35E-08 | G | A | 1800310 | 0.53 | -0.00273 | 2.58E-03 |
| rs11925699 | rs2935303 | chr3:193321115 | 0.70 | Cells - Cultured fibroblasts | A=0.47 | -0.0588 | 7.60E-05 | G | A | 1800310 | 0.53 | -0.00273 | 2.58E-03 |

| MTAG SNP | eQTL SNP | Position GRCH37 | $R^2$ | Tissue | Effect AF | Effect_Size | $P_{eQTL}$ | a1 | a2 | $N_{MTAG}$ | a1 frequency | Beta | $P_{MTAG}$ |
|---|---|---|---|---|---|---|---|---|---|---|---|---|---|
| rs11925699 | rs4597663 | chr3:193321115 | 1.00 | Whole Blood | A=0.47 | -0.1059 | 2.35E-08 | G | A | 1800310 | 0.53 | -0.00273 | 2.58E-03 |
| rs11925699 | rs4597663 | chr3:193321115 | 1.00 | Cells - Cultured fibroblasts | A=0.47 | -0.0588 | 7.60E-05 | G | A | 1800310 | 0.53 | -0.00273 | 2.58E-03 |
| rs6797738 | rs4597663 | chr3:193321173 | 0.88 | Cells - Cultured fibroblasts | A=0.439 | -0.0676 | 3.62E-06 | G | A | 1800310 | 0.56 | -0.00294 | 1.27E-03 |
| rs6797738 | rs4597663 | chr3:193321173 | 0.88 | Whole Blood | A=0.439 | -0.0973 | 2.51E-07 | G | A | 1800310 | 0.56 | -0.00294 | 1.27E-03 |
| rs9819038 | rs4597663 | chr3:193322870 | 0.88 | Cells - Cultured fibroblasts | G=0.439 | -0.0676 | 3.62E-06 | T | G | 1800310 | 0.56 | -0.00291 | 1.39E-03 |
| rs9819038 | rs4597663 | chr3:193322870 | 0.88 | Whole Blood | G=0.439 | -0.0973 | 2.51E-07 | T | G | 1800310 | 0.56 | -0.00291 | 1.39E-03 |
| rs7613578 | rs4597663 | chr3:193326458 | 0.98 | Cells - Cultured fibroblasts | T=0.475 | -0.0606 | 4.30E-05 | C | T | 1491190 | 0.53 | -0.0029 | 2.69E-03 |
| rs7613578 | rs4597663 | chr3:193326458 | 0.98 | Whole Blood | T=0.475 | -0.1036 | 4.26E-08 | C | T | 1491190 | 0.53 | -0.0029 | 2.69E-03 |
| rs7613578 | rs2935303 | chr3:193326458 | 0.68 | Cells - Cultured fibroblasts | T=0.475 | -0.0606 | 4.30E-05 | C | T | 1491190 | 0.53 | -0.0029 | 2.69E-03 |
| rs7613578 | rs2935303 | chr3:193326458 | 0.68 | Whole Blood | T=0.475 | -0.1036 | 4.26E-08 | C | T | 1491190 | 0.53 | -0.0029 | 2.69E-03 |
| rs2101340 | rs4597663 | chr3:193328895 | 0.88 | Whole Blood | G=0.439 | -0.0973 | 2.51E-07 | A | G | 1800310 | 0.56 | -0.00289 | 1.54E-03 |
| rs2101340 | rs4597663 | chr3:193328895 | 0.88 | Cells - Cultured fibroblasts | G=0.439 | -0.0676 | 3.62E-06 | A | G | 1800310 | 0.56 | -0.00289 | 1.54E-03 |
| rs13075244 | rs4597663 | chr3:193329300 | 1.00 | Cells - Cultured fibroblasts | C=0.47 | -0.0586 | 7.01E-05 | T | C | 1800310 | 0.52 | -0.00287 | 1.54E-03 |
| rs13075244 | rs4597663 | chr3:193329300 | 1.00 | Whole Blood | C=0.47 | -0.1020 | 6.64E-08 | T | C | 1800310 | 0.52 | -0.00287 | 1.54E-03 |
| rs13075244 | rs2935303 | chr3:193329300 | 0.70 | Cells - Cultured fibroblasts | C=0.47 | -0.0586 | 7.01E-05 | T | C | 1800310 | 0.52 | -0.00287 | 1.54E-03 |
| rs13075244 | rs2935303 | chr3:193329300 | 0.70 | Whole Blood | C=0.47 | -0.1020 | 6.64E-08 | T | C | 1800310 | 0.52 | -0.00287 | 1.54E-03 |
| rs13095315 | rs2935303 | chr3:193329318 | 0.70 | Whole Blood | G=0.47 | -0.1036 | 4.26E-08 | A | G | 1800310 | 0.52 | -0.00281 | 1.95E-03 |
| rs13095315 | rs4597663 | chr3:193329318 | 1.00 | Cells - Cultured fibroblasts | G=0.47 | -0.0606 | 4.30E-05 | A | G | 1800310 | 0.52 | -0.00281 | 1.95E-03 |
| rs13095315 | rs4597663 | chr3:193329318 | 1.00 | Whole Blood | G=0.47 | -0.1036 | 4.26E-08 | A | G | 1800310 | 0.52 | -0.00281 | 1.95E-03 |
| rs13095315 | rs2935303 | chr3:193329318 | 0.70 | Cells - Cultured fibroblasts | G=0.47 | -0.0606 | 4.30E-05 | A | G | 1800310 | 0.52 | -0.00281 | 1.95E-03 |
| rs9828315 | rs4597663 | chr3:193330022 | 1.00 | Whole Blood | G=0.47 | -0.1036 | 4.26E-08 | T | G | 1800310 | 0.52 | -0.0029 | 1.39E-03 |
| rs9828315 | rs2935303 | chr3:193330022 | 0.70 | Whole Blood | G=0.47 | -0.1036 | 4.26E-08 | T | G | 1800310 | 0.52 | -0.0029 | 1.39E-03 |
| rs9828315 | rs4597663 | chr3:193330022 | 1.00 | Cells - Cultured fibroblasts | G=0.47 | -0.0606 | 4.30E-05 | T | G | 1800310 | 0.52 | -0.0029 | 1.39E-03 |
| rs9828315 | rs2935303 | chr3:193330022 | 0.70 | Cells - Cultured fibroblasts | G=0.47 | -0.0606 | 4.30E-05 | T | G | 1800310 | 0.52 | -0.0029 | 1.39E-03 |
| rs6444732 | rs2935303 | chr3:193330118 | 0.70 | Whole Blood | G=0.47 | -0.1013 | 9.45E-08 | T | G | 1800310 | 0.53 | -0.00296 | 1.08E-03 |
| rs6444732 | rs2935303 | chr3:193330118 | 0.70 | Cells - Cultured fibroblasts | G=0.47 | -0.0596 | 6.73E-05 | T | G | 1800310 | 0.53 | -0.00296 | 1.08E-03 |
| rs6444732 | rs4597663 | chr3:193330118 | 1.00 | Cells - Cultured fibroblasts | G=0.47 | -0.0596 | 6.73E-05 | T | G | 1800310 | 0.53 | -0.00296 | 1.08E-03 |
| rs6444732 | rs4597663 | chr3:193330118 | 1.00 | Whole Blood | G=0.47 | -0.1013 | 9.45E-08 | T | G | 1800310 | 0.53 | -0.00296 | 1.08E-03 |
| rs7646539 | rs4597663 | chr3:193334846 | 0.88 | Whole Blood | G=0.439 | -0.0982 | 1.83E-07 | A | G | 1800310 | 0.56 | -0.00287 | 1.67E-03 |
| rs7646539 | rs4597663 | chr3:193334846 | 0.88 | Cells - Cultured fibroblasts | G=0.439 | -0.0671 | 4.13E-06 | A | G | 1800310 | 0.56 | -0.00287 | 1.67E-03 |

| MTAG SNP | eQTL SNP | Position GRCH37 | $R^2$ | Tissue | Effect AF | Effect_Size | $P_{eQTL}$ | a1 | a2 | $N_{MTAG}$ | a1 frequency | Beta | $P_{MTAG}$ |
|---|---|---|---|---|---|---|---|---|---|---|---|---|---|
| rs7624750 | rs4597663 | chr3:193334991 | 1.00 | Whole Blood | A=0.47 | -0.1045 | 3.17E-08 | G | A | 1800310 | 0.52 | -0.00294 | 1.18E-03 |
| rs7624750 | rs2935303 | chr3:193334991 | 0.70 | Whole Blood | A=0.47 | -0.1045 | 3.17E-08 | G | A | 1800310 | 0.52 | -0.00294 | 1.18E-03 |
| rs7624750 | rs4597663 | chr3:193334991 | 1.00 | Cells - Cultured fibroblasts | A=0.47 | -0.0606 | 4.30E-05 | G | A | 1800310 | 0.52 | -0.00294 | 1.18E-03 |
| rs7624750 | rs2935303 | chr3:193334991 | 0.70 | Cells - Cultured fibroblasts | A=0.47 | -0.0606 | 4.30E-05 | G | A | 1800310 | 0.52 | -0.00294 | 1.18E-03 |
| rs10937593 | rs4597663 | chr3:193335252 | 1.00 | Whole Blood | T=0.47 | -0.1045 | 3.17E-08 | G | T | 1800310 | 0.52 | -0.0029 | 1.38E-03 |
| rs10937593 | rs4597663 | chr3:193335252 | 1.00 | Cells - Cultured fibroblasts | T=0.47 | -0.0606 | 4.30E-05 | G | T | 1800310 | 0.52 | -0.0029 | 1.38E-03 |
| rs10937593 | rs2935303 | chr3:193335252 | 0.70 | Whole Blood | T=0.47 | -0.1045 | 3.17E-08 | G | T | 1800310 | 0.52 | -0.0029 | 1.38E-03 |
| rs10937593 | rs2935303 | chr3:193335252 | 0.70 | Cells - Cultured fibroblasts | T=0.47 | -0.0606 | 4.30E-05 | G | T | 1800310 | 0.52 | -0.0029 | 1.38E-03 |
| rs9832709 | rs4597663 | chr3:193336425 | 0.88 | Cells - Cultured fibroblasts | C=0.439 | -0.0676 | 3.62E-06 | T | C | 1800310 | 0.56 | -0.0029 | 1.47E-03 |
| rs9832709 | rs4597663 | chr3:193336425 | 0.88 | Whole Blood | C=0.439 | -0.0982 | 1.91E-07 | T | C | 1800310 | 0.56 | -0.0029 | 1.47E-03 |
| rs3772393 | rs4597663 | chr3:193336639 | 0.88 | Whole Blood | C=0.439 | -0.0982 | 1.91E-07 | T | C | 1701420 | 0.56 | -0.0035 | 1.81E-04 |
| rs3772393 | rs4597663 | chr3:193336639 | 0.88 | Cells - Cultured fibroblasts | C=0.439 | -0.0676 | 3.62E-06 | T | C | 1701420 | 0.56 | -0.0035 | 1.81E-04 |
| rs9817704 | rs4597663 | chr3:193337021 | 1.00 | Cells - Cultured fibroblasts | A=0.47 | -0.0606 | 4.30E-05 | G | A | 1800310 | 0.52 | -0.00289 | 1.41E-03 |
| rs9817704 | rs2935303 | chr3:193337021 | 0.70 | Cells - Cultured fibroblasts | A=0.47 | -0.0606 | 4.30E-05 | G | A | 1800310 | 0.52 | -0.00289 | 1.41E-03 |
| rs9817704 | rs2935303 | chr3:193337021 | 0.70 | Whole Blood | A=0.47 | -0.1045 | 3.17E-08 | G | A | 1800310 | 0.52 | -0.00289 | 1.41E-03 |
| rs9817704 | rs4597663 | chr3:193337021 | 1.00 | Whole Blood | A=0.47 | -0.1045 | 3.17E-08 | G | A | 1800310 | 0.52 | -0.00289 | 1.41E-03 |
| rs6770913 | rs2935303 | chr3:193338335 | 0.70 | Whole Blood | A=0.47 | -0.1036 | 4.26E-08 | C | A | 1800310 | 0.52 | -0.00288 | 1.46E-03 |
| rs6770913 | rs4597663 | chr3:193338335 | 1.00 | Whole Blood | A=0.47 | -0.1036 | 4.26E-08 | C | A | 1800310 | 0.52 | -0.00288 | 1.46E-03 |
| rs6770913 | rs4597663 | chr3:193338335 | 1.00 | Cells - Cultured fibroblasts | A=0.47 | -0.0606 | 4.30E-05 | C | A | 1800310 | 0.52 | -0.00288 | 1.46E-03 |
| rs6770913 | rs2935303 | chr3:193338335 | 0.70 | Cells - Cultured fibroblasts | A=0.47 | -0.0606 | 4.30E-05 | C | A | 1800310 | 0.52 | -0.00288 | 1.46E-03 |
| rs34520293 | rs4597663 | chr3:193340310 | 1.00 | Whole Blood | C=0.47 | -0.1036 | 4.26E-08 | A | C | 1701420 | 0.53 | -0.00356 | 1.27E-04 |
| rs34520293 | rs2935303 | chr3:193340310 | 0.70 | Whole Blood | C=0.47 | -0.1036 | 4.26E-08 | A | C | 1701420 | 0.53 | -0.00356 | 1.27E-04 |
| rs34520293 | rs4597663 | chr3:193340310 | 1.00 | Cells - Cultured fibroblasts | C=0.47 | -0.0606 | 4.30E-05 | A | C | 1701420 | 0.53 | -0.00356 | 1.27E-04 |
| rs34520293 | rs2935303 | chr3:193340310 | 0.70 | Cells - Cultured fibroblasts | C=0.47 | -0.0606 | 4.30E-05 | A | C | 1701420 | 0.53 | -0.00356 | 1.27E-04 |
| rs12490391 | rs4597663 | chr3:193341016 | 0.88 | Cells - Cultured fibroblasts | T=0.439 | -0.0676 | 3.62E-06 | C | T | 1800310 | 0.56 | -0.00286 | 1.70E-03 |
| rs12490391 | rs4597663 | chr3:193341016 | 0.88 | Whole Blood | T=0.439 | -0.0973 | 2.51E-07 | C | T | 1800310 | 0.56 | -0.00286 | 1.70E-03 |
| rs6444733 | rs2935303 | chr3:193346648 | 0.70 | Whole Blood | C=0.47 | -0.1010 | 8.37E-08 | A | C | 1701420 | 0.53 | -0.00358 | 1.19E-04 |
| rs6444733 | rs4597663 | chr3:193346648 | 1.00 | Whole Blood | C=0.47 | -0.1010 | 8.37E-08 | A | C | 1701420 | 0.53 | -0.00358 | 1.19E-04 |
| rs6444734 | rs4597663 | chr3:193346921 | 0.88 | Cells - Cultured fibroblasts | T=0.439 | -0.0676 | 3.62E-06 | C | T | 1445630 | 0.53 | -0.00352 | 4.49E-04 |
| rs6444734 | rs4597663 | chr3:193346921 | 0.88 | Whole Blood | T=0.439 | -0.0973 | 2.51E-07 | C | T | 1445630 | 0.53 | -0.00352 | 4.49E-04 |

| MTAG SNP | eQTL SNP | Position GRCH37 | $R^2$ | Tissue | Effect AF | Effect_Size | $P_{eQTL}$ | a1 | a2 | $N_{MTAG}$ | a1 frequency | Beta | $P_{MTAG}$ |
|---|---|---|---|---|---|---|---|---|---|---|---|---|---|
| rs12696671 | rs4597663 | chr3:193348919 | 0.88 | Cells - Cultured fibroblasts | G=0.439 | -0.0676 | 3.62E-06 | A | G | 1800310 | 0.56 | -0.00286 | 1.74E-03 |
| rs12696671 | rs4597663 | chr3:193348919 | 0.88 | Whole Blood | G=0.439 | -0.0973 | 2.51E-07 | A | G | 1800310 | 0.56 | -0.00286 | 1.74E-03 |
| rs2101341 | rs2935303 | chr3:193350294 | 0.70 | Cells - Cultured fibroblasts | C=0.47 | -0.0589 | 6.94E-05 | T | C | 1800310 | 0.52 | -0.00291 | 1.34E-03 |
| rs2101341 | rs2935303 | chr3:193350294 | 0.70 | Whole Blood | C=0.47 | -0.1059 | 1.83E-08 | T | C | 1800310 | 0.52 | -0.00291 | 1.34E-03 |
| rs2101341 | rs4597663 | chr3:193350294 | 1.00 | Whole Blood | C=0.47 | -0.1059 | 1.83E-08 | T | C | 1800310 | 0.52 | -0.00291 | 1.34E-03 |
| rs2101341 | rs4597663 | chr3:193350294 | 1.00 | Cells - Cultured fibroblasts | C=0.47 | -0.0589 | 6.94E-05 | T | C | 1800310 | 0.52 | -0.00291 | 1.34E-03 |
| rs13096511 | rs4597663 | chr3:193350837 | 0.87 | Cells - Cultured fibroblasts | G=0.434 | -0.0676 | 3.62E-06 | T | G | 1723250 | 0.56 | -0.00304 | 1.01E-03 |
| rs13096511 | rs4597663 | chr3:193350837 | 0.87 | Whole Blood | G=0.434 | -0.0973 | 2.51E-07 | T | G | 1723250 | 0.56 | -0.00304 | 1.01E-03 |
| rs13078398 | rs4597663 | chr3:193350876 | 0.88 | Whole Blood | T=0.439 | -0.0973 | 2.51E-07 | G | T | 1800310 | 0.56 | -0.00285 | 1.78E-03 |
| rs13078398 | rs4597663 | chr3:193350876 | 0.88 | Cells - Cultured fibroblasts | T=0.439 | -0.0676 | 3.62E-06 | G | T | 1800310 | 0.56 | -0.00285 | 1.78E-03 |
| rs3736198 | rs2935303 | chr3:193353072 | 0.70 | Whole Blood | A=0.47 | -0.1036 | 4.26E-08 | G | A | 1701420 | 0.52 | -0.00365 | 8.65E-05 |
| rs3736198 | rs4597663 | chr3:193353072 | 1.00 | Whole Blood | A=0.47 | -0.1036 | 4.26E-08 | G | A | 1701420 | 0.52 | -0.00365 | 8.65E-05 |
| rs3736198 | rs2935303 | chr3:193353072 | 0.70 | Cells - Cultured fibroblasts | A=0.47 | -0.0606 | 4.30E-05 | G | A | 1701420 | 0.52 | -0.00365 | 8.65E-05 |
| rs3736198 | rs4597663 | chr3:193353072 | 1.00 | Cells - Cultured fibroblasts | A=0.47 | -0.0606 | 4.30E-05 | G | A | 1701420 | 0.52 | -0.00365 | 8.65E-05 |
| rs6807435 | rs4597663 | chr3:193353955 | 1.00 | Cells - Cultured fibroblasts | A=0.47 | -0.0606 | 4.30E-05 | G | A | 1800310 | 0.53 | -0.0029 | 1.40E-03 |
| rs6807435 | rs2935303 | chr3:193353955 | 0.70 | Cells - Cultured fibroblasts | A=0.47 | -0.0606 | 4.30E-05 | G | A | 1800310 | 0.53 | -0.0029 | 1.40E-03 |
| rs6807435 | rs2935303 | chr3:193353955 | 0.70 | Whole Blood | A=0.47 | -0.1047 | 2.84E-08 | G | A | 1800310 | 0.53 | -0.0029 | 1.40E-03 |
| rs6807435 | rs4597663 | chr3:193353955 | 1.00 | Whole Blood | A=0.47 | -0.1047 | 2.84E-08 | G | A | 1800310 | 0.53 | -0.0029 | 1.40E-03 |
| rs78386430 | rs78912308 | chr3:193356770 | 0.94 | Cells - Cultured fibroblasts | G=0.096 | -0.2416 | 5.34E-22 | A | G | 1522690 | 0.90 | -0.00556 | 6.27E-04 |
| rs78386430 | rs78912308 | chr3:193356770 | 0.94 | Whole Blood | G=0.096 | -0.1830 | 9.05E-07 | A | G | 1522690 | 0.90 | -0.00556 | 6.27E-04 |
| rs1078932 | rs4597663 | chr3:193358672 | 0.88 | Cells - Cultured fibroblasts | A=0.439 | -0.0676 | 3.62E-06 | G | A | 1800310 | 0.56 | -0.00282 | 1.96E-03 |
| rs1078932 | rs4597663 | chr3:193358672 | 0.88 | Whole Blood | A=0.439 | -0.0973 | 2.51E-07 | G | A | 1800310 | 0.56 | -0.00282 | 1.96E-03 |
| rs36146795 | rs4597663 | chr3:193359193 | 0.87 | Whole Blood | C=0.434 | -0.0973 | 2.51E-07 | T | C | 1723250 | 0.56 | -0.00303 | 1.08E-03 |
| rs36146795 | rs4597663 | chr3:193359193 | 0.87 | Cells - Cultured fibroblasts | C=0.434 | -0.0699 | 1.63E-06 | T | C | 1723250 | 0.56 | -0.00303 | 1.08E-03 |
| rs7630464 | rs1872532 | chr3:193359565 | 0.93 | Cells - Cultured fibroblasts | G=0.313 | -0.0864 | 3.48E-08 | A | G | 1800310 | 0.67 | -0.00293 | 2.36E-03 |
| rs7630464 | rs1872532 | chr3:193359565 | 0.93 | Whole Blood | G=0.313 | -0.0933 | 6.51E-06 | A | G | 1800310 | 0.67 | -0.00293 | 2.36E-03 |
| rs1016226 | rs4597663 | chr3:193361617 | 0.87 | Cells - Cultured fibroblasts | T=0.434 | -0.0680 | 3.04E-06 | C | T | 1800310 | 0.56 | -0.00269 | 3.24E-03 |
| rs1016226 | rs4597663 | chr3:193361617 | 0.87 | Whole Blood | T=0.434 | -0.0985 | 1.88E-07 | C | T | 1800310 | 0.56 | -0.00269 | 3.24E-03 |
| rs77303706 | rs78912308 | chr3:193361917 | 0.94 | Cells - Cultured fibroblasts | A=0.096 | -0.2416 | 5.34E-22 | G | A | 1621580 | 0.90 | -0.00451 | 4.64E-03 |
| rs77303706 | rs78912308 | chr3:193361917 | 0.94 | Whole Blood | A=0.096 | -0.1830 | 9.05E-07 | G | A | 1621580 | 0.90 | -0.00451 | 4.64E-03 |

| MTAG SNP | eQTL SNP | Position GRCH37 | $R^2$ | Tissue | Effect AF | Effect_Size | $P_{eQTL}$ | a1 | a2 | $N_{MTAG}$ | a1 frequency | Beta | $P_{MTAG}$ |
|---|---|---|---|---|---|---|---|---|---|---|---|---|---|
| rs11916478 | rs4597663 | chr3:193362189 | 0.87 | Whole Blood | A=0.434 | -0.0957 | 4.08E-07 | G | A | 1800310 | 0.56 | -0.00275 | 2.56E-03 |
| rs11916478 | rs4597663 | chr3:193362189 | 0.87 | Cells - Cultured fibroblasts | A=0.434 | -0.0703 | 1.34E-06 | G | A | 1800310 | 0.56 | -0.00275 | 2.56E-03 |
| rs6771174 | rs1872532 | chr3:193362666 | 0.81 | Whole Blood | T=0.283 | -0.0871 | 4.21E-05 | C | T | 1800310 | 0.71 | -0.00299 | 2.69E-03 |
| rs6771174 | rs1872532 | chr3:193362666 | 0.81 | Cells - Cultured fibroblasts | T=0.283 | -0.0972 | 5.74E-10 | C | T | 1800310 | 0.71 | -0.00299 | 2.69E-03 |
| rs2291373 | rs4597663 | chr3:193363163 | 0.96 | Cells - Cultured fibroblasts | A=0.47 | -0.0596 | 5.91E-05 | G | A | 1800310 | 0.53 | -0.00275 | 2.45E-03 |
| rs2291373 | rs2935303 | chr3:193363163 | 0.73 | Cells - Cultured fibroblasts | A=0.47 | -0.0596 | 5.91E-05 | G | A | 1800310 | 0.53 | -0.00275 | 2.45E-03 |
| rs2291373 | rs4597663 | chr3:193363163 | 0.96 | Whole Blood | A=0.47 | -0.0979 | 2.52E-07 | G | A | 1800310 | 0.53 | -0.00275 | 2.45E-03 |
| rs2291373 | rs2935303 | chr3:193363163 | 0.73 | Whole Blood | A=0.47 | -0.0979 | 2.52E-07 | G | A | 1800310 | 0.53 | -0.00275 | 2.45E-03 |
| rs77539393 | rs78912308 | chr3:193363267 | 0.94 | Whole Blood | C=0.096 | -0.1838 | 4.66E-07 | T | C | 1621580 | 0.90 | -0.0045 | 4.69E-03 |
| rs77539393 | rs78912308 | chr3:193363267 | 0.94 | Cells - Cultured fibroblasts | C=0.096 | -0.2426 | 6.82E-23 | T | C | 1621580 | 0.90 | -0.0045 | 4.69E-03 |
| rs7621979 | rs4597663 | chr3:193364483 | 0.86 | Whole Blood | T=0.444 | -0.0952 | 4.86E-07 | G | T | 1645020 | 0.56 | -0.00295 | 1.68E-03 |
| rs7621979 | rs4597663 | chr3:193364483 | 0.86 | Cells - Cultured fibroblasts | T=0.444 | -0.0665 | 5.23E-06 | G | T | 1645020 | 0.56 | -0.00295 | 1.68E-03 |
| rs9845217 | rs4597663 | chr3:193365544 | 0.98 | Whole Blood | G=0.475 | -0.0973 | 2.76E-07 | A | G | 1800310 | 0.53 | -0.00286 | 1.61E-03 |
| rs9845217 | rs4597663 | chr3:193365544 | 0.98 | Cells - Cultured fibroblasts | G=0.475 | -0.0576 | 1.07E-04 | A | G | 1800310 | 0.53 | -0.00286 | 1.61E-03 |
| rs9845217 | rs2935303 | chr3:193365544 | 0.71 | Whole Blood | G=0.475 | -0.0973 | 2.76E-07 | A | G | 1800310 | 0.53 | -0.00286 | 1.61E-03 |
| rs9845217 | rs2935303 | chr3:193365544 | 0.71 | Cells - Cultured fibroblasts | G=0.475 | -0.0576 | 1.07E-04 | A | G | 1800310 | 0.53 | -0.00286 | 1.61E-03 |
| rs9831900 | rs2935303 | chr3:193365974 | 0.71 | Cells - Cultured fibroblasts | G=0.475 | -0.0610 | 3.76E-05 | T | G | 1213560 | 0.49 | -0.00356 | 8.12E-04 |
| rs9831900 | rs4597663 | chr3:193365974 | 0.98 | Cells - Cultured fibroblasts | G=0.475 | -0.0610 | 3.76E-05 | T | G | 1213560 | 0.49 | -0.00356 | 8.12E-04 |
| rs9831900 | rs2935303 | chr3:193365974 | 0.71 | Whole Blood | G=0.475 | -0.1034 | 4.24E-08 | T | G | 1213560 | 0.49 | -0.00356 | 8.12E-04 |
| rs9831900 | rs4597663 | chr3:193365974 | 0.98 | Whole Blood | G=0.475 | -0.1034 | 4.24E-08 | T | G | 1213560 | 0.49 | -0.00356 | 8.12E-04 |
| rs7620116 | rs4597663 | chr3:193368692 | 0.85 | Cells - Cultured fibroblasts | A=0.439 | -0.0669 | 4.39E-06 | C | A | 1701420 | 0.56 | -0.00333 | 3.69E-04 |
| rs7620116 | rs4597663 | chr3:193368692 | 0.85 | Whole Blood | A=0.439 | -0.0964 | 3.67E-07 | C | A | 1701420 | 0.56 | -0.00333 | 3.69E-04 |
| rs7620272 | rs4597663 | chr3:193368700 | 0.85 | Whole Blood | A=0.439 | -0.0964 | 3.67E-07 | G | A | 1800310 | 0.56 | -0.00271 | 3.01E-03 |
| rs7620272 | rs4597663 | chr3:193368700 | 0.85 | Cells - Cultured fibroblasts | A=0.439 | -0.0669 | 4.39E-06 | G | A | 1800310 | 0.56 | -0.00271 | 3.01E-03 |
| rs11717605 | rs4597663 | chr3:193369463 | 0.85 | Whole Blood | G=0.439 | -0.0952 | 4.86E-07 | A | G | 1800310 | 0.56 | -0.00286 | 1.74E-03 |
| rs11717605 | rs4597663 | chr3:193369463 | 0.85 | Cells - Cultured fibroblasts | G=0.439 | -0.0669 | 4.39E-06 | A | G | 1800310 | 0.56 | -0.00286 | 1.74E-03 |
| rs56263480 | rs78912308 | chr3:193372027 | 0.94 | Whole Blood | A=0.096 | -0.1849 | 4.79E-07 | G | A | 1621580 | 0.90 | -0.00449 | 4.87E-03 |
| rs56263480 | rs78912308 | chr3:193372027 | 0.94 | Cells - Cultured fibroblasts | A=0.096 | -0.2407 | 2.90E-22 | G | A | 1621580 | 0.90 | -0.00449 | 4.87E-03 |
| rs56263480 | rs78912308 | chr3:193372027 | 0.94 | Pancreas | A=0.096 | -0.3050 | 3.79E-05 | G | A | 1621580 | 0.90 | -0.00449 | 4.87E-03 |
| rs79129003 | rs78912308 | chr3:193373936 | 0.94 | Pancreas | A=0.096 | -0.3050 | 3.79E-05 | G | A | 1621580 | 0.90 | -0.00444 | 5.30E-03 |

| MTAG SNP | eQTL SNP | Position GRCH37 | $R^2$ | Tissue | Effect AF | Effect_Size | $P_{eQTL}$ | a1 | a2 | $N_{MTAG}$ | a1 frequency | Beta | $P_{MTAG}$ |
|---|---|---|---|---|---|---|---|---|---|---|---|---|---|
| rs79129003 | rs78912308 | chr3:193373936 | 0.94 | Cells - Cultured fibroblasts | A=0.096 | -0.2407 | 2.90E-22 | G | A | 1621580 | 0.90 | -0.00444 | 5.30E-03 |
| rs79129003 | rs78912308 | chr3:193373936 | 0.94 | Whole Blood | A=0.096 | -0.1849 | 4.79E-07 | G | A | 1621580 | 0.90 | -0.00444 | 5.30E-03 |
| rs6444735 | rs2935303 | chr3:193374384 | 0.73 | Whole Blood | T=0.47 | -0.1046 | 3.15E-08 | C | T | 1701420 | 0.53 | -0.00345 | 2.10E-04 |
| rs6444735 | rs4597663 | chr3:193374384 | 0.96 | Whole Blood | T=0.47 | -0.1046 | 3.15E-08 | C | T | 1701420 | 0.53 | -0.00345 | 2.10E-04 |
| rs6444735 | rs4597663 | chr3:193374384 | 0.96 | Cells - Cultured fibroblasts | T=0.47 | -0.0614 | 3.21E-05 | C | T | 1701420 | 0.53 | -0.00345 | 2.10E-04 |
| rs6444735 | rs2935303 | chr3:193374384 | 0.73 | Cells - Cultured fibroblasts | T=0.47 | -0.0614 | 3.21E-05 | C | T | 1701420 | 0.53 | -0.00345 | 2.10E-04 |
| rs9851685 | rs2935303 | chr3:193374964 | 0.71 | Cells - Cultured fibroblasts | C=0.475 | -0.0610 | 3.76E-05 | T | C | 1800310 | 0.53 | -0.00273 | 2.64E-03 |
| rs9851685 | rs4597663 | chr3:193374964 | 0.98 | Whole Blood | C=0.475 | -0.1046 | 3.15E-08 | T | C | 1800310 | 0.53 | -0.00273 | 2.64E-03 |
| rs9851685 | rs4597663 | chr3:193374964 | 0.98 | Cells - Cultured fibroblasts | C=0.475 | -0.0610 | 3.76E-05 | T | C | 1800310 | 0.53 | -0.00273 | 2.64E-03 |
| rs9851685 | rs2935303 | chr3:193374964 | 0.71 | Whole Blood | C=0.475 | -0.1046 | 3.15E-08 | T | C | 1800310 | 0.53 | -0.00273 | 2.64E-03 |
| rs9837255 | rs4597663 | chr3:193376227 | 0.85 | Cells - Cultured fibroblasts | T=0.439 | -0.0669 | 4.39E-06 | C | T | 1645020 | 0.56 | -0.00281 | 2.78E-03 |
| rs9837255 | rs4597663 | chr3:193376227 | 0.85 | Whole Blood | T=0.439 | -0.0964 | 3.67E-07 | C | T | 1645020 | 0.56 | -0.00281 | 2.78E-03 |
| rs9837682 | rs4597663 | chr3:193376298 | 0.85 | Whole Blood | A=0.439 | -0.0964 | 3.67E-07 | G | A | 1800310 | 0.56 | -0.0027 | 3.06E-03 |
| rs9837682 | rs4597663 | chr3:193376298 | 0.85 | Cells - Cultured fibroblasts | A=0.439 | -0.0669 | 4.39E-06 | G | A | 1800310 | 0.56 | -0.0027 | 3.06E-03 |
| rs28457829 | rs4597663 | chr3:193377037 | 0.85 | Cells - Cultured fibroblasts | A=0.439 | -0.0669 | 4.39E-06 | G | A | 1701420 | 0.56 | -0.00333 | 3.73E-04 |
| rs28457829 | rs4597663 | chr3:193377037 | 0.85 | Whole Blood | A=0.439 | -0.0964 | 3.67E-07 | G | A | 1701420 | 0.56 | -0.00333 | 3.73E-04 |
| rs9291060 | rs4597663 | chr3:193378126 | 0.85 | Cells - Cultured fibroblasts | C=0.439 | -0.0669 | 4.39E-06 | A | C | 1701420 | 0.56 | -0.00332 | 3.88E-04 |
| rs9291060 | rs4597663 | chr3:193378126 | 0.85 | Whole Blood | C=0.439 | -0.0964 | 3.67E-07 | A | C | 1701420 | 0.56 | -0.00332 | 3.88E-04 |
| rs9813526 | rs4597663 | chr3:193381915 | 0.85 | Whole Blood | C=0.439 | -0.0957 | 3.90E-07 | T | C | 1800310 | 0.56 | -0.00281 | 2.04E-03 |
| rs9813526 | rs4597663 | chr3:193381915 | 0.85 | Cells - Cultured fibroblasts | C=0.439 | -0.0669 | 4.39E-06 | T | C | 1800310 | 0.56 | -0.00281 | 2.04E-03 |
| rs9831353 | rs4597663 | chr3:193382138 | 0.85 | Whole Blood | G=0.439 | -0.1000 | 8.78E-08 | A | G | 1800310 | 0.56 | -0.00269 | 3.18E-03 |
| rs9831353 | rs4597663 | chr3:193382138 | 0.85 | Cells - Cultured fibroblasts | G=0.439 | -0.0670 | 3.23E-06 | A | G | 1800310 | 0.56 | -0.00269 | 3.18E-03 |
| rs9869814 | rs2935303 | chr3:193382370 | 0.73 | Whole Blood | A=0.47 | -0.1039 | 3.38E-08 | G | A | 1800310 | 0.52 | -0.00279 | 2.07E-03 |
| rs9869814 | rs4597663 | chr3:193382370 | 0.96 | Cells - Cultured fibroblasts | A=0.47 | -0.0614 | 3.21E-05 | G | A | 1800310 | 0.52 | -0.00279 | 2.07E-03 |
| rs9869814 | rs2935303 | chr3:193382370 | 0.73 | Cells - Cultured fibroblasts | A=0.47 | -0.0614 | 3.21E-05 | G | A | 1800310 | 0.52 | -0.00279 | 2.07E-03 |
| rs9869814 | rs4597663 | chr3:193382370 | 0.96 | Whole Blood | A=0.47 | -0.1039 | 3.38E-08 | G | A | 1800310 | 0.52 | -0.00279 | 2.07E-03 |
| rs10937594 | rs4597663 | chr3:193383943 | 0.85 | Whole Blood | C=0.439 | -0.1007 | 8.13E-08 | T | C | 1800310 | 0.56 | -0.00256 | 4.92E-03 |
| rs10937594 | rs4597663 | chr3:193383943 | 0.85 | Cells - Cultured fibroblasts | C=0.439 | -0.0670 | 3.23E-06 | T | C | 1800310 | 0.56 | -0.00256 | 4.92E-03 |
| rs10804957 | rs2935303 | chr3:193383973 | 0.73 | Whole Blood | G=0.47 | -0.1046 | 3.15E-08 | A | G | 1701420 | 0.53 | -0.00341 | 2.48E-04 |
| rs10804957 | rs2935303 | chr3:193383973 | 0.73 | Cells - Cultured fibroblasts | G=0.47 | -0.0614 | 3.21E-05 | A | G | 1701420 | 0.53 | -0.00341 | 2.48E-04 |

| MTAG SNP | eQTL SNP | Position GRCH37 | $R^2$ | Tissue | Effect AF | Effect_Size | $P_{eQTL}$ | a1 | a2 | $N_{MTAG}$ | a1 frequency | Beta | $P_{MTAG}$ |
|---|---|---|---|---|---|---|---|---|---|---|---|---|---|
| rs10804957 | rs4597663 | chr3:193383973 | 0.96 | Cells - Cultured fibroblasts | G=0.47 | -0.0614 | 3.21E-05 | A | G | 1701420 | 0.53 | -0.00341 | 2.48E-04 |
| rs10804957 | rs4597663 | chr3:193383973 | 0.96 | Whole Blood | G=0.47 | -0.1046 | 3.15E-08 | A | G | 1701420 | 0.53 | -0.00341 | 2.48E-04 |
| rs13070791 | rs4597663 | chr3:193385275 | 0.83 | Whole Blood | T=0.434 | -0.0969 | 3.30E-07 | C | T | 1800310 | 0.57 | -0.00284 | 1.85E-03 |
| rs13070791 | rs4597663 | chr3:193385275 | 0.83 | Cells - Cultured fibroblasts | T=0.434 | -0.0667 | 4.45E-06 | C | T | 1800310 | 0.57 | -0.00284 | 1.85E-03 |
| rs12494482 | rs4597663 | chr3:193386239 | 0.96 | Cells - Cultured fibroblasts | T=0.47 | -0.0614 | 3.21E-05 | C | T | 1800310 | 0.53 | -0.00276 | 2.37E-03 |
| rs12494482 | rs4597663 | chr3:193386239 | 0.96 | Whole Blood | T=0.47 | -0.1046 | 3.15E-08 | C | T | 1800310 | 0.53 | -0.00276 | 2.37E-03 |
| rs12494482 | rs2935303 | chr3:193386239 | 0.73 | Whole Blood | T=0.47 | -0.1046 | 3.15E-08 | C | T | 1800310 | 0.53 | -0.00276 | 2.37E-03 |
| rs12494482 | rs2935303 | chr3:193386239 | 0.73 | Cells - Cultured fibroblasts | T=0.47 | -0.0614 | 3.21E-05 | C | T | 1800310 | 0.53 | -0.00276 | 2.37E-03 |
| rs7616448 | rs4597663 | chr3:193387345 | 0.85 | Whole Blood | A=0.439 | -0.0964 | 3.67E-07 | G | A | 1800310 | 0.56 | -0.00265 | 3.65E-03 |
| rs7616448 | rs4597663 | chr3:193387345 | 0.85 | Cells - Cultured fibroblasts | A=0.439 | -0.0669 | 4.39E-06 | G | A | 1800310 | 0.56 | -0.00265 | 3.65E-03 |
| rs11915238 | rs4597663 | chr3:193387831 | 0.96 | Whole Blood | C=0.47 | -0.1038 | 4.35E-08 | T | C | 1800310 | 0.53 | -0.00281 | 1.94E-03 |
| rs11915238 | rs4597663 | chr3:193387831 | 0.96 | Cells - Cultured fibroblasts | C=0.47 | -0.0610 | 3.43E-05 | T | C | 1800310 | 0.53 | -0.00281 | 1.94E-03 |
| rs11915238 | rs2935303 | chr3:193387831 | 0.73 | Cells - Cultured fibroblasts | C=0.47 | -0.0610 | 3.43E-05 | T | C | 1800310 | 0.53 | -0.00281 | 1.94E-03 |
| rs11915238 | rs2935303 | chr3:193387831 | 0.73 | Whole Blood | C=0.47 | -0.1038 | 4.35E-08 | T | C | 1800310 | 0.53 | -0.00281 | 1.94E-03 |
| rs10937596 | rs4597663 | chr3:193387937 | 0.96 | Whole Blood | A=0.47 | -0.1046 | 3.15E-08 | C | A | 1800310 | 0.53 | -0.00275 | 2.46E-03 |
| rs10937596 | rs4597663 | chr3:193387937 | 0.96 | Cells - Cultured fibroblasts | A=0.47 | -0.0614 | 3.21E-05 | C | A | 1800310 | 0.53 | -0.00275 | 2.46E-03 |
| rs10937596 | rs2935303 | chr3:193387937 | 0.73 | Whole Blood | A=0.47 | -0.1046 | 3.15E-08 | C | A | 1800310 | 0.53 | -0.00275 | 2.46E-03 |
| rs10937596 | rs2935303 | chr3:193387937 | 0.73 | Cells - Cultured fibroblasts | A=0.47 | -0.0614 | 3.21E-05 | C | A | 1800310 | 0.53 | -0.00275 | 2.46E-03 |
| rs1985005 | rs2935303 | chr3:193388237 | 0.73 | Whole Blood | T=0.47 | -0.1041 | 3.46E-08 | C | T | 1800310 | 0.53 | -0.00268 | 3.17E-03 |
| rs1985005 | rs2935303 | chr3:193388237 | 0.73 | Cells - Cultured fibroblasts | T=0.47 | -0.0609 | 3.41E-05 | C | T | 1800310 | 0.53 | -0.00268 | 3.17E-03 |
| rs1985005 | rs4597663 | chr3:193388237 | 0.96 | Cells - Cultured fibroblasts | T=0.47 | -0.0609 | 3.41E-05 | C | T | 1800310 | 0.53 | -0.00268 | 3.17E-03 |
| rs1985005 | rs4597663 | chr3:193388237 | 0.96 | Whole Blood | T=0.47 | -0.1041 | 3.46E-08 | C | T | 1800310 | 0.53 | -0.00268 | 3.17E-03 |
| rs11720294 | rs4597663 | chr3:193389390 | 0.96 | Cells - Cultured fibroblasts | A=0.47 | -0.0609 | 3.41E-05 | G | A | 1800310 | 0.53 | -0.00269 | 3.02E-03 |
| rs11720294 | rs4597663 | chr3:193389390 | 0.96 | Whole Blood | A=0.47 | -0.1041 | 3.46E-08 | G | A | 1800310 | 0.53 | -0.00269 | 3.02E-03 |
| rs11720294 | rs2935303 | chr3:193389390 | 0.73 | Cells - Cultured fibroblasts | A=0.47 | -0.0609 | 3.41E-05 | G | A | 1800310 | 0.53 | -0.00269 | 3.02E-03 |
| rs11720294 | rs2935303 | chr3:193389390 | 0.73 | Whole Blood | A=0.47 | -0.1041 | 3.46E-08 | G | A | 1800310 | 0.53 | -0.00269 | 3.02E-03 |
| rs11720234 | rs4597663 | chr3:193389413 | 0.85 | Whole Blood | A=0.439 | -0.0975 | 2.91E-07 | C | A | 1701420 | 0.56 | -0.00331 | 4.01E-04 |
| rs11720234 | rs4597663 | chr3:193389413 | 0.85 | Cells - Cultured fibroblasts | A=0.439 | -0.0683 | 2.65E-06 | C | A | 1701420 | 0.56 | -0.00331 | 4.01E-04 |
| rs11720340 | rs2935303 | chr3:193389485 | 0.73 | Whole Blood | A=0.47 | -0.1030 | 4.94E-08 | G | A | 1645020 | 0.53 | -0.00275 | 3.28E-03 |
| rs11720340 | rs4597663 | chr3:193389485 | 0.96 | Cells - Cultured fibroblasts | A=0.47 | -0.0594 | 5.25E-05 | G | A | 1645020 | 0.53 | -0.00275 | 3.28E-03 |

| MTAG SNP | eQTL SNP | Position GRCH37 | $R^2$ | Tissue | Effect AF | Effect_Size | $P_{eQTL}$ | a1 | a2 | $N_{MTAG}$ | a1 frequency | Beta | $P_{MTAG}$ |
|---|---|---|---|---|---|---|---|---|---|---|---|---|---|
| rs11720340 | rs2935303 | chr3:193389485 | 0.73 | Cells - Cultured fibroblasts | A=0.47 | -0.0594 | 5.25E-05 | G | A | 1645020 | 0.53 | -0.00275 | 3.28E-03 |
| rs11720340 | rs4597663 | chr3:193389485 | 0.96 | Whole Blood | A=0.47 | -0.1030 | 4.94E-08 | G | A | 1645020 | 0.53 | -0.00275 | 3.28E-03 |
| rs6796713 | rs2935303 | chr3:193391085 | 0.73 | Whole Blood | A=0.47 | -0.1019 | 6.62E-08 | G | A | 1800310 | 0.52 | -0.0028 | 2.00E-03 |
| rs6796713 | rs4597663 | chr3:193391085 | 0.96 | Cells - Cultured fibroblasts | A=0.47 | -0.0594 | 5.25E-05 | G | A | 1800310 | 0.52 | -0.0028 | 2.00E-03 |
| rs6796713 | rs4597663 | chr3:193391085 | 0.96 | Whole Blood | A=0.47 | -0.1019 | 6.62E-08 | G | A | 1800310 | 0.52 | -0.0028 | 2.00E-03 |
| rs6796713 | rs2935303 | chr3:193391085 | 0.73 | Cells - Cultured fibroblasts | A=0.47 | -0.0594 | 5.25E-05 | G | A | 1800310 | 0.52 | -0.0028 | 2.00E-03 |
| rs6444737 | rs4597663 | chr3:193391680 | 0.96 | Whole Blood | G=0.47 | -0.1034 | 4.24E-08 | A | G | 1800310 | 0.52 | -0.00285 | 1.67E-03 |
| rs6444737 | rs2935303 | chr3:193391680 | 0.73 | Cells - Cultured fibroblasts | G=0.47 | -0.0618 | 2.69E-05 | A | G | 1800310 | 0.52 | -0.00285 | 1.67E-03 |
| rs6444737 | rs2935303 | chr3:193391680 | 0.73 | Whole Blood | G=0.47 | -0.1034 | 4.24E-08 | A | G | 1800310 | 0.52 | -0.00285 | 1.67E-03 |
| rs6444737 | rs4597663 | chr3:193391680 | 0.96 | Cells - Cultured fibroblasts | G=0.47 | -0.0618 | 2.69E-05 | A | G | 1800310 | 0.52 | -0.00285 | 1.67E-03 |
| rs78898827 | rs78912308 | chr3:193392197 | 0.94 | Whole Blood | G=0.096 | -0.1830 | 9.05E-07 | A | G | 1621580 | 0.90 | -0.0047 | 3.11E-03 |
| rs78898827 | rs78912308 | chr3:193392197 | 0.94 | Cells - Cultured fibroblasts | G=0.096 | -0.2416 | 5.34E-22 | A | G | 1621580 | 0.90 | -0.0047 | 3.11E-03 |
| rs3851984 | rs2935303 | chr3:193393612 | 0.73 | Cells - Cultured fibroblasts | G=0.47 | -0.0614 | 3.21E-05 | A | G | 1800310 | 0.53 | -0.00274 | 2.48E-03 |
| rs3851984 | rs4597663 | chr3:193393612 | 0.96 | Whole Blood | G=0.47 | -0.1046 | 3.15E-08 | A | G | 1800310 | 0.53 | -0.00274 | 2.48E-03 |
| rs3851984 | rs4597663 | chr3:193393612 | 0.96 | Cells - Cultured fibroblasts | G=0.47 | -0.0614 | 3.21E-05 | A | G | 1800310 | 0.53 | -0.00274 | 2.48E-03 |
| rs3851984 | rs2935303 | chr3:193393612 | 0.73 | Whole Blood | G=0.47 | -0.1046 | 3.15E-08 | A | G | 1800310 | 0.53 | -0.00274 | 2.48E-03 |
| rs4443116 | rs4597663 | chr3:193401021 | 0.96 | Whole Blood | G=0.47 | -0.0983 | 1.75E-07 | A | G | 1800310 | 0.52 | -0.00284 | 1.75E-03 |
| rs4443116 | rs2935303 | chr3:193401021 | 0.73 | Whole Blood | G=0.47 | -0.0983 | 1.75E-07 | A | G | 1800310 | 0.52 | -0.00284 | 1.75E-03 |
| rs1806741 | rs4597663 | chr3:193401874 | 0.85 | Whole Blood | T=0.439 | -0.0958 | 4.02E-07 | C | T | 1800310 | 0.56 | -0.00261 | 4.28E-03 |
| rs1806741 | rs4597663 | chr3:193401874 | 0.85 | Cells - Cultured fibroblasts | T=0.439 | -0.0655 | 6.83E-06 | C | T | 1800310 | 0.56 | -0.00261 | 4.28E-03 |
| rs2367721 | rs4597663 | chr3:193402003 | 0.85 | Cells - Cultured fibroblasts | G=0.439 | -0.0635 | 1.30E-05 | A | G | 1800310 | 0.56 | -0.00267 | 3.40E-03 |
| rs2367721 | rs4597663 | chr3:193402003 | 0.85 | Whole Blood | G=0.439 | -0.0948 | 5.36E-07 | A | G | 1800310 | 0.56 | -0.00267 | 3.40E-03 |
| rs885053 | rs4597663 | chr3:193402008 | 0.85 | Whole Blood | C=0.439 | -0.1007 | 8.13E-08 | T | C | 1701420 | 0.56 | -0.00316 | 7.30E-04 |
| rs885053 | rs4597663 | chr3:193402008 | 0.85 | Cells - Cultured fibroblasts | C=0.439 | -0.0670 | 3.23E-06 | T | C | 1701420 | 0.56 | -0.00316 | 7.30E-04 |
| rs6444739 | rs4597663 | chr3:193403048 | 0.85 | Cells - Cultured fibroblasts | A=0.439 | -0.0669 | 4.39E-06 | G | A | 1800310 | 0.56 | -0.00283 | 1.93E-03 |
| rs6444739 | rs4597663 | chr3:193403048 | 0.85 | Whole Blood | A=0.439 | -0.0952 | 4.86E-07 | G | A | 1800310 | 0.56 | -0.00283 | 1.93E-03 |
| rs79911093 | rs78912308 | chr3:193405301 | 0.94 | Whole Blood | A=0.096 | -0.1849 | 4.79E-07 | G | A | 1621580 | 0.90 | -0.00461 | 3.77E-03 |
| rs79911093 | rs78912308 | chr3:193405301 | 0.94 | Cells - Cultured fibroblasts | A=0.096 | -0.2407 | 2.90E-22 | G | A | 1621580 | 0.90 | -0.00461 | 3.77E-03 |
| rs79911093 | rs78912308 | chr3:193405301 | 0.94 | Pancreas | A=0.096 | -0.3050 | 3.79E-05 | G | A | 1621580 | 0.90 | -0.00461 | 3.77E-03 |
| rs12696674 | rs4597663 | chr3:193407447 | 0.85 | Cells - Cultured fibroblasts | T=0.439 | -0.0669 | 4.39E-06 | C | T | 1800310 | 0.56 | -0.00281 | 2.03E-03 |

| MTAG SNP | eQTL SNP | Position GRCH37 | $R^2$ | Tissue | Effect AF | Effect_Size | $P_{eQTL}$ | a1 | a2 | $N_{MTAG}$ | a1 frequency | Beta | $P_{MTAG}$ |
|---|---|---|---|---|---|---|---|---|---|---|---|---|---|
| rs12696674 | rs4597663 | chr3:193407447 | 0.85 | Whole Blood | T=0.439 | -0.0952 | 4.86E-07 | C | T | 1800310 | 0.56 | -0.00281 | 2.03E-03 |
| rs13068219 | rs4597663 | chr3:193407650 | 0.85 | Whole Blood | A=0.439 | -0.0958 | 4.02E-07 | G | A | 1800310 | 0.56 | -0.00263 | 3.96E-03 |
| rs13068219 | rs4597663 | chr3:193407650 | 0.85 | Cells - Cultured fibroblasts | A=0.439 | -0.0655 | 6.83E-06 | G | A | 1800310 | 0.56 | -0.00263 | 3.96E-03 |
| rs13067959 | rs4597663 | chr3:193407703 | 0.85 | Whole Blood | A=0.439 | -0.0955 | 4.58E-07 | C | A | 1800310 | 0.56 | -0.00286 | 1.72E-03 |
| rs13067959 | rs4597663 | chr3:193407703 | 0.85 | Cells - Cultured fibroblasts | A=0.439 | -0.0673 | 4.14E-06 | C | A | 1800310 | 0.56 | -0.00286 | 1.72E-03 |
| rs6793945 | rs4597663 | chr3:193407889 | 0.85 | Whole Blood | G=0.439 | -0.0995 | 1.19E-07 | A | G | 1800310 | 0.55 | -0.00271 | 2.92E-03 |
| rs6793945 | rs4597663 | chr3:193407889 | 0.85 | Cells - Cultured fibroblasts | G=0.439 | -0.0672 | 3.36E-06 | A | G | 1800310 | 0.55 | -0.00271 | 2.92E-03 |
| rs7638928 | rs4597663 | chr3:193408419 | 0.85 | Whole Blood | G=0.439 | -0.0952 | 4.86E-07 | A | G | 1800310 | 0.56 | -0.00282 | 1.95E-03 |
| rs7638928 | rs4597663 | chr3:193408419 | 0.85 | Cells - Cultured fibroblasts | G=0.439 | -0.0669 | 4.39E-06 | A | G | 1800310 | 0.56 | -0.00282 | 1.95E-03 |
| rs111999715 | rs78912308 | chr3:193411322 | 0.89 | Pancreas | T=0.091 | -0.3368 | 1.14E-05 | G | T | 1445630 | 0.90 | -0.00552 | 7.71E-04 |
| rs111999715 | rs78912308 | chr3:193411322 | 0.89 | Whole Blood | T=0.091 | -0.1853 | 7.34E-07 | G | T | 1445630 | 0.90 | -0.00552 | 7.71E-04 |
| rs111999715 | rs78912308 | chr3:193411322 | 0.89 | Cells - Cultured fibroblasts | T=0.091 | -0.2406 | 2.08E-21 | G | T | 1445630 | 0.90 | -0.00552 | 7.71E-04 |
| rs1056390 | rs4597663 | chr3:193414481 | 0.85 | Whole Blood | T=0.439 | -0.0956 | 3.86E-07 | C | T | 1701420 | 0.55 | -0.00322 | 5.58E-04 |
| rs1056390 | rs4597663 | chr3:193414481 | 0.85 | Cells - Cultured fibroblasts | T=0.439 | -0.0634 | 1.23E-05 | C | T | 1701420 | 0.55 | -0.00322 | 5.58E-04 |
| rs11719309 | rs4597663 | chr3:193414780 | 0.85 | Whole Blood | C=0.439 | -0.0956 | 3.86E-07 | T | C | 1645020 | 0.55 | -0.00277 | 3.16E-03 |
| rs11719309 | rs4597663 | chr3:193414780 | 0.85 | Cells - Cultured fibroblasts | C=0.439 | -0.0634 | 1.23E-05 | T | C | 1645020 | 0.55 | -0.00277 | 3.16E-03 |
| rs56329083 | rs78912308 | chr3:193415416 | 0.89 | Whole Blood | G=0.091 | -0.1873 | 3.85E-07 | A | G | 1621580 | 0.90 | -0.00459 | 3.86E-03 |
| rs56329083 | rs78912308 | chr3:193415416 | 0.89 | Pancreas | G=0.091 | -0.3404 | 6.44E-06 | A | G | 1621580 | 0.90 | -0.00459 | 3.86E-03 |
| rs56329083 | rs78912308 | chr3:193415416 | 0.89 | Cells - Cultured fibroblasts | G=0.091 | -0.2401 | 1.02E-21 | A | G | 1621580 | 0.90 | -0.00459 | 3.86E-03 |
| rs12152411 | rs4597663 | chr3:193415679 | 0.85 | Whole Blood | C=0.439 | -0.0956 | 3.86E-07 | T | C | 1800310 | 0.55 | -0.00265 | 3.55E-03 |
| rs12152411 | rs4597663 | chr3:193415679 | 0.85 | Cells - Cultured fibroblasts | C=0.439 | -0.0634 | 1.23E-05 | T | C | 1800310 | 0.55 | -0.00265 | 3.55E-03 |
| rs77918412 | rs78912308 | chr3:193416448 | 0.89 | Pancreas | T=0.091 | -0.3404 | 6.44E-06 | C | T | 1621580 | 0.90 | -0.00465 | 3.39E-03 |
| rs77918412 | rs78912308 | chr3:193416448 | 0.89 | Cells - Cultured fibroblasts | T=0.091 | -0.2401 | 1.02E-21 | C | T | 1621580 | 0.90 | -0.00465 | 3.39E-03 |
| rs77918412 | rs78912308 | chr3:193416448 | 0.89 | Whole Blood | T=0.091 | -0.1873 | 3.85E-07 | C | T | 1621580 | 0.90 | -0.00465 | 3.39E-03 |
| rs7652334 | rs4597663 | chr3:193420857 | 0.85 | Whole Blood | C=0.439 | -0.0896 | 2.50E-06 | T | C | 1800310 | 0.55 | -0.00269 | 3.16E-03 |
| rs7652334 | rs4597663 | chr3:193420857 | 0.85 | Cells - Cultured fibroblasts | C=0.439 | -0.0605 | 4.22E-05 | T | C | 1800310 | 0.55 | -0.00269 | 3.16E-03 |
| rs6787463 | rs2935303 | chr3:193318291 | 0.66 | Whole Blood | A=0.49 | -0.1020 | 6.22E-08 | G | A | 1800310 | 0.52 | -0.00275 | 2.37E-03 |
| rs6787463 | rs4597663 | chr3:193318291 | 0.92 | Cells - Cultured fibroblasts | A=0.49 | -0.0569 | 1.06E-04 | G | A | 1800310 | 0.52 | -0.00275 | 2.37E-03 |
| rs6787463 | rs2935303 | chr3:193318291 | 0.66 | Cells - Cultured fibroblasts | A=0.49 | -0.0569 | 1.06E-04 | G | A | 1800310 | 0.52 | -0.00275 | 2.37E-03 |
| rs6787463 | rs4597663 | chr3:193318291 | 0.92 | Whole Blood | A=0.49 | -0.1020 | 6.22E-08 | G | A | 1800310 | 0.52 | -0.00275 | 2.37E-03 |

| MTAG SNP | eQTL SNP | Position GRCH37 | $R^2$ | Tissue | Effect AF | Effect_Size | $P_{eQTL}$ | a1 | a2 | $N_{MTAG}$ | a1 frequency | Beta | $P_{MTAG}$ |
|---|---|---|---|---|---|---|---|---|---|---|---|---|---|
| rs6764269 | rs6764269 | chr3:193422395 | 1.00 | Whole Blood | C=0.591 | -0.0857 | 4.52E-06 | T | C | 1800310 | 0.46 | -0.00265 | 3.48E-03 |

**Table S5. Summary of key published human evidence for OPA1 expression in MASLD and liver cirrhosis.**

| Source | Human tissue/cohort | Assay | Key OPA1 finding |
|---|---|---|---|
| Human Protein Atlas[6, 7] | Normal human tissues | RNA-seq / IHC | Moderate hepatic OPA1 protein staining and relatively high OPA1 RNA expression. |
| Da Dalt et al., 2023[8] | GSE135251 + GSE207310 | Bulk RNA-seq reanalysis | Higher hepatic OPA1 mRNA; positive association with NAFLD Activity Score in both cohorts. |
| Moore et al., 2022[9] | n=130; control/NAFLD/borderline NASH/definite NASH | Western blot | Hepatic OPA1 protein decreased from NAFLD onset through NASH progression. |
| Niu et al., 2022[10] | Control n=15; NASH n=20; cirrhosis n=10 | DIA-MS proteomics | No significant total OPA1 difference in NASH or cirrhosis versus control after adjusted analysis. |
| Zhao et al., 2013[11] | HCC/cirrhotic/non-cirrhotic liver tissues | Western blot | Indirect suggestion of lower OPA1 by the cirrhotic stage; no direct cirrhosis-versus-normal estimate. |
